# Height, but not binding epitope, affects the potency of synthetic TCR agonists

**DOI:** 10.1101/2021.05.13.443919

**Authors:** Kiera B. Wilhelm, Shumpei Morita, Darren B. McAffee, Sungi Kim, Mark K. O’Dair, Jay T. Groves

## Abstract

Under physiological conditions, peptide-MHC (pMHC) molecules can trigger T-cell receptors (TCRs) as monovalent ligands that are sparsely distributed on the plasma membrane of an antigen-presenting cell. TCR can also be activated by artificial clustering, such as with pMHC tetramers or antibodies; however, these strategies circumvent many of the natural ligand discrimination mechanisms of the T cell and can elicit non-physiological signaling activity. We have recently introduced a synthetic TCR agonist composed of an anti-TCRβ Fab’ antibody fragment covalently bound to a DNA oligonucleotide, which serves as a membrane anchor. This Fab’-DNA ligand efficiently activates TCR as a monomer when membrane-associated and exhibits a potency and activation profile resembling agonist pMHC. In this report, we explore the geometric requirements for effective TCR triggering and cellular activation by Fab’-DNA ligands. We find that T cells are insensitive to the ligand binding epitope on the TCR complex, but that length of the DNA tether is important. Increasing the intermembrane distance spanned by Fab’-DNA:TCR complexes decreases TCR triggering efficiency and T cell activation potency, consistent with the kinetic-segregation model of TCR triggering. These results establish design parameters for construction of synthetic TCR agonists that are able to activate polyclonal T cell populations, such as T cells from a human patient, in a similar manner as the native pMHC ligand.

**STATEMENT OF SIGNIFICANCE:** We report geometric requirements for potent T cell activation by synthetic TCR ligands that mimic biophysical properties of the native pMHC ligand, but have the additional ability to activate polyclonal T cell populations. We find that increasing the space between apposed membranes at TCR binding events decreases ligand potency, but that changing the ligand’s binding epitope on the TCR has essentially no effect. The observed decrease in potency with increased ligand height is attributed to the longer ligands’ attenuated ability to trigger TCR at binding events.

## INTRODUCTION

T cells play a central role in adaptive immunity by recognizing foreign peptide fragments presented in MHC molecules (pMHC) with their T-cell receptors (TCR). In order to identify a wide range of potential pathogenic peptides, each individual develops a polyclonal repertoire of T cells with distinct TCR genes. In developing thymocytes, the region of the TCR gene that encodes the pMHC recognition site undergoes somatic recombination, which creates a large sample space of potential receptors, and cells with TCR clonotypes that successfully pass a screening process survive (1). The resulting diversity of TCRs within an individual is critical for successfully conferring adaptive immunity, but presents challenges to the study of T cell activation because the cognate pMHC that activates a given T cell is generally not known.

Antibodies that bind the TCR and associated CD3 complex (TCR/CD3) readily activate T cells, as measured by cytokine secretion, proliferation, and changes in surface receptor expression (2–6), but this method of stimulation differs from physiological antigen activation. Anti-TCR/CD3 antibodies, as well as widely used pMHC tetramers (7), induce crosslinking of TCR on the T cell surface. This crosslinking is an essential aspect of their activation mechanism; neither pMHC monomer nor monovalent Fab’ antibody fragments are active from solution (7–10), and anti-TCR/CD3 antibodies in solution must typically be further crosslinked by secondary antibodies for full activity (11). By contrast, a growing body of evidence indicates that membrane-associated pMHC molecules are highly active as monomers (12–15). Furthermore, at physiological densities of agonist pMHC (0.1-2 µm^-2^) (16–18), only tens of individual pMHC:TCR ligation events, which are widely spaced within the cell interface, are sufficient to activate T cells (19, 20). At higher agonist pMHC densities (∼10-250 µm^-2^), pMHC:TCR complexes form microclusters (21, 22) that further reorganize into the large scale pattern of the immunological synapse (23, 24), but these larger scale organizations at high antigen density are not required for T cell activation (12–15, 20).

We have recently developed a class of synthetic TCR agonists composed of an anti-TCR/CD3 Fab’ fragment covalently bound to a DNA oligonucleotide that can uniformly activate polyclonal T cell populations as membrane-associated monomers (25). Like natively monomeric pMHC, Fab’-DNA molecules are inactive from solution at any concentration, but are highly potent when conjugated to a supported lipid bilayer (SLB) via DNA hybridization. Previous work focused on the anti-TCRβ H57-597 Fab’-DNA (H57 Fab’-DNA) and found it to exhibit a similar potency as a strong pMHC agonist (25). Mechanistic studies of TCR triggering suggest that the Fab’ binding epitope on the TCR/CD3 complex (26, 27) and/or the intermembrane spacing established at TCR binding events may impact ligand potency (28–32).

Here we examine a panel of Fab’-DNA constructs that bind three distinct epitopes on the TCR/CD3 complex and have DNA tethers of various lengths (Fig. 1). We synthesized Fab’-DNA ligands derived from the anti-CD3ε 145-2C11 antibody (2C11 Fab’-DNA) and the anti-CD3ε/γ 17A2 antibody (17A2 Fab’-DNA), in addition to the previously reported anti-TCRβ H57 Fab’-DNA (3–5, 26, 33) (Fig. 1B). Both 2C11 and H57 bivalent antibodies are commonly used to activate T cells and bind at or near the reportedly mechanosensitive FG loop of TCRβ (26, 27, 33–35). The 17A2 antibody is less frequently used to activate T cells and has been reported to be less potent due to its binding geometry to the TCR/CD3 complex (26, 27). DNA tethers ranging from 16 to 76 nucleotides were designed for all Fabs, creating 14 to 50 nm space between apposed membranes at binding events (Fig. 1C). This range of sizes spans from the native pMHC:TCR intermembrane spacing of about 14 nm (36–38) to beyond the spacing of about 30 nm established by ICAM-1:LFA-1 adhesion complexes (39) and the large 21-40 nm extracellular domain of CD45 (32, 40, 41), which has long been thought to be sterically excluded by the short pMHC:TCR complex size at T cell – antigen presenting cell junctions (21, 28–30).

**Figure 1.**
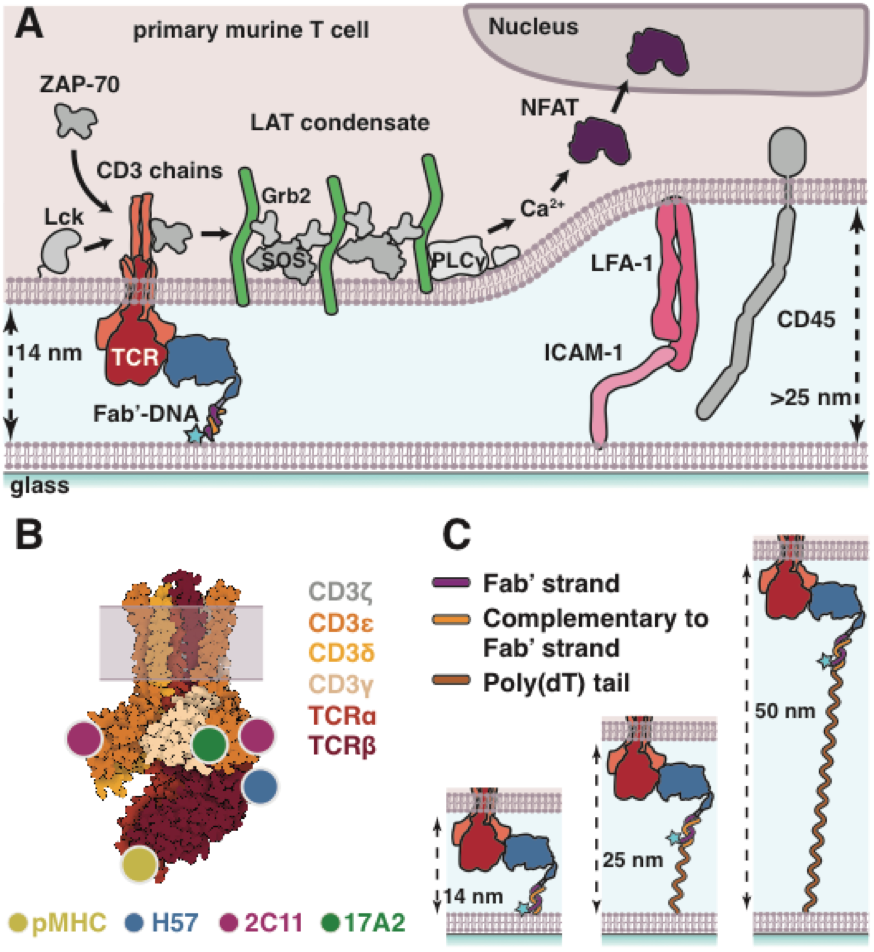
Fab’-DNA is a modular TCR ligand. (**A**) Fab’-DNA and ICAM-1 are presented on supported lipid bilayers. Upon adding T cells to the bilayer, LFA-1 adheres to ICAM-1 to create a stable intermembrane junction between the SLB and T cell. Within this junction, Fab’-DNA binds TCR, creating a narrow 14 nm space between membranes that excludes ICAM-1:LFA-1 conjugates and the phosphatase CD45 from close proximity. Productive TCR triggering is monitored by the formation of TCR-proximal LAT condensates and cellular activation is monitored by translocation of the transcription factor NFAT from the cytosol to the nucleus. (**B**) Fab’-DNA constructs are synthesized from antibodies that bind three distinct epitopes on the TCR/CD3 complex (adapted from PDB entry 6JXR). Binding epitopes for pMHC and all Fabs have previously been determined by crystal and NMR structures and are approximately mapped onto the TCR/CD3 cryoEM structure. (**C**) The spacing between the SLB and T cell at Fab’-DNA:TCR binding events is modulated by varying the length of the DNA tether. A poly(dT) tail is added to the membrane-proximal 5’ end of the thiol-DNA, increasing the allowed distance between the membrane and the region of the thiol-DNA to which Fab’-DNA anneals.

We find that the intermembrane space established by DNA tether length dramatically affects Fab’-DNA potency, with larger spacing leading to less potent cellular activation, but that varying the Fab’ binding epitope has essentially no effect. T cell activation was monitored by imaging translocation of a fluorescent reporter for the nuclear factor for the activation of T cells (NFAT) from the cytosol to the nucleus, which provides a binary readout of successful activation of the calcium signaling pathway in T cells (19, 20, 25, 42). TCR-proximal signaling, measured by localized formation of protein condensates of linker for the activation of T cells (LAT) also showed the same trend. LAT is an intrinsically disordered protein that serves a signaling scaffold immediately downstream of TCR triggering (43, 44) and its condensate formation proximal to individual Fab’-DNA:TCR complexes is an indicator of local signaling activity (45, 46). The observed dependence of T cell signaling on Fab’-DNA height is consistent with the kinetic-segregation model of TCR triggering, which implicates steric exclusion of bulky phosphatases from closely apposed membrane regions at pMHC:TCR binding events as a driving force for signal propagation from the TCR (28–32). These results are also consistent with studies of similarly-structured bispecific T cell engagers (BiTEs) that show BiTEs are more effective if they bind epitopes creating smaller intermembrane spaces where they bridge TCRs on a T cell to melanoma markers on an opposing cell (47, 48). The T cells’ indifferent response to ligand binding epitope suggests that any intramolecular aspects of the TCR triggering mechanism (27, 35, 49) do not strongly depend on TCR engagement geometry in the context of adhesion molecules and a cell-cell interface.

## MATERIALS AND METHODS

Detailed descriptions for all subsections below can be found in the Supporting Materials.

### Fab’-DNA synthesis

All antibodies were digested with pepsin or Glu-C endoproteinase to retain the cysteines in the hinge region and then partially reduced with 2-mercaptoethylamine to form Fab’ fragments. Maleimide-functionalized, dye-labeled DNA oligonucleotides were then conjugated to the reduced cysteine residue on the Fab’ to form crude Fab’-DNA. This product was purified by size exclusion and anion exchange chromatography to obtain monomeric Fab’ conjugated to a single DNA oligonucleotide that was labeled with a single fluorophore (25).

### Supported Lipid Bilayer Preparation

Low-defect, fluid supported lipid bilayers (SLB) were prepared from a solution of small unilamellar vesicles (SUVs). SUVs were first prepared by mixing 95% DOPC (1,2-dioleoyl-sn-glycero-3-phosphocholine), 3% MCC-DOPE (1,2-dioleoyl-sn-glycero-3-phosphoethanolamine-N-[4-(p-maleimidomethyl)cyclohexane-carboxamide] sodium salt), and 2% Ni-NTA-DOGS (1,2-dioleoyl-sn-glycero-3-[(N-(5-amino-1-carboxypentyl)iminodiacetic acid)succinyl] nickel salt) phospholipids in chloroform in a piranha-etched flask, drying and resuspending lipids in Milli-Q water to 0.5 mg/mL total lipid concentration, and sonicating with a probe sonicator. The SUVs were then centrifuged to remove titanium particles and lipid aggregates and mixed 1:1 with 1x PBS, resulting in a spreading solution with a final concentration of 0.25 mg/mL total lipid content. Supported membranes were formed by vesicle fusion of SUVs by adding this spreading solution to imaging chambers assembled with freshly piranha-etched glass coverslips. Thiol-DNA, complementary to the Fab’-DNA strand, was deprotected with 10 mM TCEP for 90 min to expose free thiol and then incubated on the bilayer at about 1 µM in PBS to obtain a density of approximately hundreds of molecules/µm^2^ on the SLB (50). These bilayers were stable overnight at 4 °C. Before experiments, the SLB was charged with 30 mM NiCl_2_ to ensure stable chelation of polyhistidine-tagged ICAM-1 to the NTA-DOGS lipids. Proteins to be coupled to the bilayer were prepared in imaging buffer, added to the imaging chamber, and incubated for 35 min. Fab’-DNA incubation concentrations ranged from 50 pM for single molecule studies to 10 nM for high density samples. ICAM-1 was incubated at a concentration of 100 nM.

### T cell harvesting and culture

CD4+ T cells expressing the AND TCR (51) were harvested, cultured, and transduced as previously described (15, 52). Briefly, T cells were harvested from hemizygous transgenic mice from the cross of (B10.Cg-Tg(TcrAND)53Hed/J) x (B10.BR-H2k2 H2-T18a/SgSnJ) strains (The Jackson Laboratory, Bar Harbor, ME) and activated by moth cytochrome c peptide (MCC_88-103_) immediately after harvest. IL-2 was added the following day within 24h of the harvest. Cells prepared for live cell assays were retrovirally transduced with NFAT-,mCherry, LAT-eGFP, or a LAT-eGFP P2A NFAT-mCherry plasmid-containing supertatants collected from Platinum-Eco cells (Cell Biolabs, San Diego, CA). T cells were imaged on days 5-8. Cell health was verified each day that data were collected by assessing cell morphology and signaling in response to bilayers with no agonist ligand and high density of pMHC. All animal work was approved by Lawrence Berkeley National Laboratory Animal Welfare and Research Committee under the approved protocols #17702 and #17703.

### Microscopy

Imaging experiments were performed on an inverted Nikon Eclipse Ti-E motorized inverted microscope (Nikon, Tokyo, Japan) with total internal reflection fluorescence (TIRF) microscopy, reflection interference contrast microscopy (RICM), and epifluorescence capabilities and laser lines at 405, 488, 532, and 640 nm. Freely diffusing, single Fab’-DNA-Atto647 molecules were imaged in TIRF using 20 ms exposure time, 8.6 mW power at sample, and 1000 gain. Single binding events between Fab’-DNA and TCR were imaged using 500 ms exposure time, 0.4 mW power at sample, and 1000 gain. RICM and epifluorescence images were acquired with moderate exposure time (∼100 ms) with gain depending on the intensity of the signal. LAT was imaged using low power (0.4-0.8 mW at sample), moderate exposure time (50-200 ms) and 1000 gain, with exact parameters depending on the expression of LAT-eGFP. Micro-Manager was used to automate acquisitions and collect data (53).

### Image analysis

Single particle images of freely diffusing Fab’-DNA (Fig. 2) and Fab’-DNA bound to TCR (Fig. 3) were localized and tracked in the ImageJ plugin TrackMate (54). Particles were identified using the difference of Gaussians detector, particle diameter and threshold were determined by eye, and all data from a given experiment were analyzed uniformly. The diameter was usually set to about 0.4 µm. The simple linear assignment problem (LAP) tracking algorithm was used to link localized spots. Maximum particle linking distances were set depending on the time lapse between images and particle speed. Data from single particle tracking were then exported to MATLAB. Step size distributions, step photobleaching, dwell time distributions, and fraction bound were analyzed using custom MATLAB scripts, described in detail in the Supporting Materials and Methods..

**Figure 2.**
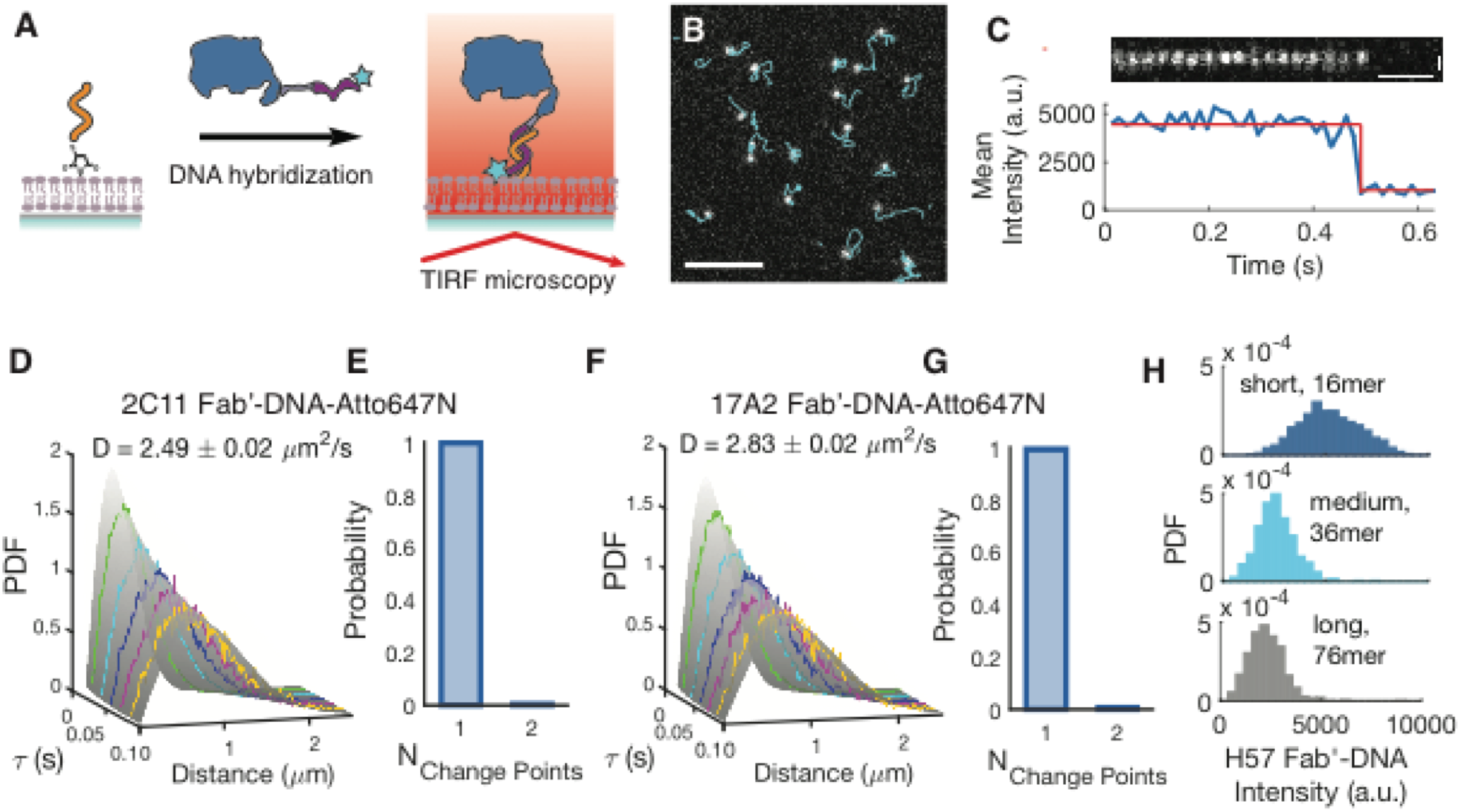
All Fab’-DNA constructs diffuse as monomers on supported lipid bilayers. (**A**) The thiol-DNA strand is covalently bound to the SLB via thiol/maleimide chemistry. Fab’-DNA that is labeled with a single Atto647N fluorophore is then incubated in the imaging chambers and anneals to the thiol-DNA strand. Fab’-DNA molecules are visualized using TIRF microscopy. (**B**) 2C11 Fab’-DNA molecules of uniform brightness undergo Brownian diffusion on the SLB. Scale bar 5 µm. (**C**) Diffusing 2C11 Fab’-DNA particles bleach in a single step, confirming that each particle is a single Fab’-DNA molecule. Vertical scale bar 500 nm. Horizontal scale bar 0.1 s. (**D** and **F**) The step size distribution of 2C11 (D) and 17A2 (F) Fab’-DNA fits a single component diffusion model well. A multi-delay time protocol was used to build step size distributions at multiple time delays and all distributions were simultaneously fit to obtain the diffusion coefficient for each ligand. Error indicates the 95% CI. (n > 50,000 steps) Data are representative of three experiments. (**E** and **G**) >99% of 2C11 (n = 122) (E) and 17A2 (n = 175) (G) Fab’-DNA particles bleach in a single step, with intensity traces exhibiting a single change point as shown in (C). These data were obtained on gel-phase SLBs to enable tracking of particles for their full trajectories until photobleaching. The <1% of particles that undergo 2-step photobleaching match the probability that two Fab’-DNA are randomly spaced below the diffraction limit. No particles bleach in three or more steps. Panels B-G display data collected with the shortest thiol-DNA. (**H**) Single particle fluorescence intensity distributions for Fab’-DNA conjugated to the SLB by annealing to short, medium, and long thiol-DNA strands show decreased fluorescence intensity with increased tether length. Short thiol-DNA: n > 1000 particles; medium thiol-DNA: n > 500 particles; long thiol-DNA: n > 1000 particles.

**Figure 3.**
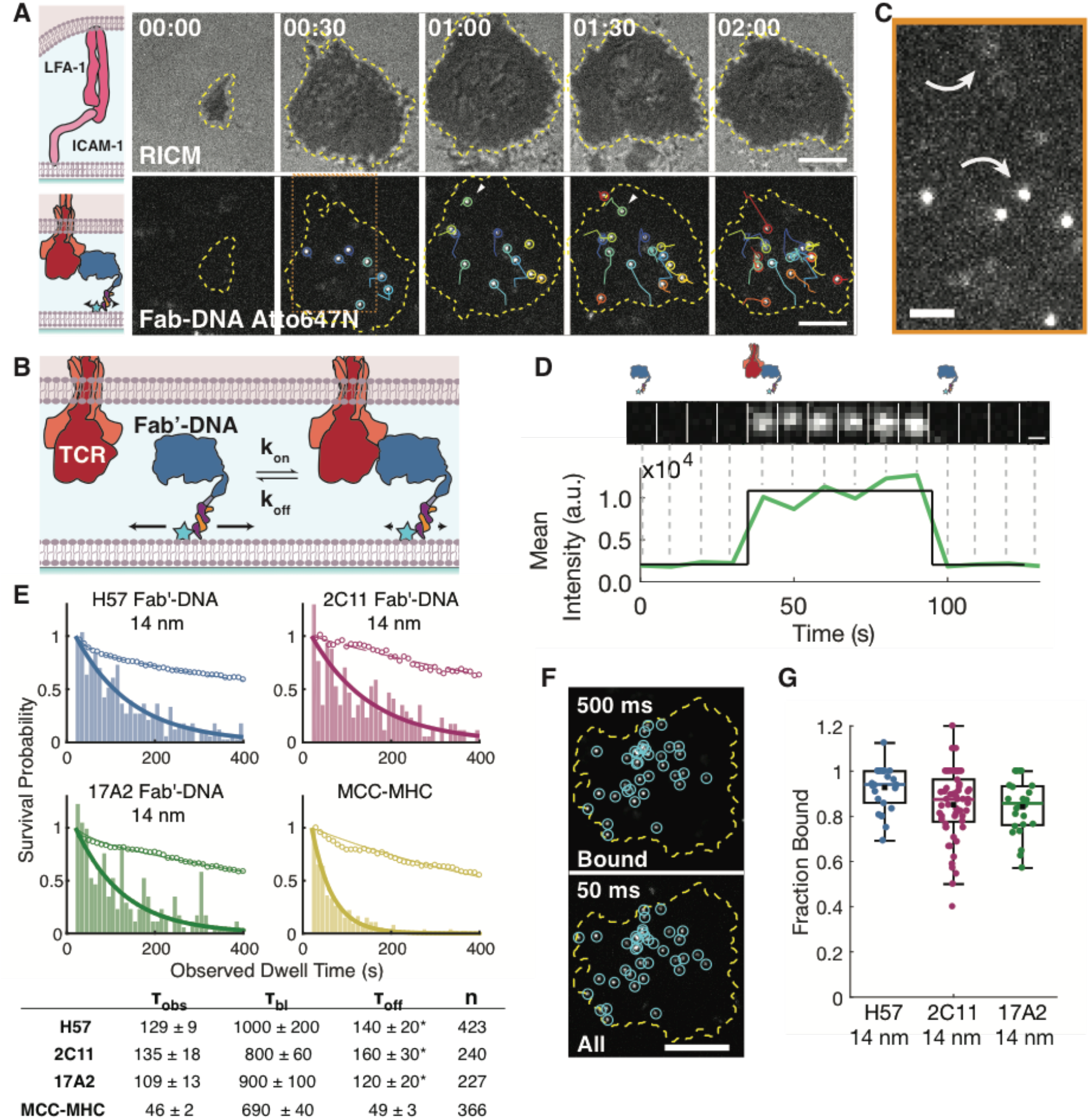
Measurements of individual binding events between Fab’-DNA and TCR illustrate that all Fab’-DNA complexes strongly bind TCR. (**A**) T cells adhere to the bilayer through ICAM-1:LFA-1 interactions (cell footprint visualized by RICM), and Fab’-DNA-Atto647N binds TCR within this SLB - T cell junction (visualized by TIRF). Each binding event is tracked through time, denoted with a unique color for each event. Scale bar 5 µm. (**B**) This schematic illustrates how bound Fab’-DNA can be distinguished from unbound Fab’-DNA by the dramatic decrease in Fab’-DNA mobility upon TCR binding. Relative mobility is represented by horizontal arrows. (**C**) An imaging strategy using a long, 500 ms exposure time at low power allows slowly-moving bound Fab’-DNA images (bottom arrow) to be resolved while quickly diffusing free Fab’-DNA forms a blurred image (top arrow). This area corresponds to the area in (A) outlined by a dotted orange line. Scale bar 2 µm. (**D**) The intensity trace of a Fab’-DNA :TCR binding event through time (white arrow in (A)) illustrates that single binding and unbinding events are readily visualized. Scale bar 300 nm. (**E**) Dwell time distributions, corrected for photobleaching (empty circles), for Fab’-DNA ligands illustrate their very slow off-rates. Fab’-DNA ligands bind for at least twice as long as the strong agonist pMHC. Data are compiled from cells from at least two mice for each condition in dwell time and fraction bound measurements. *The reported τ_off_ for Fab’-DNA ligands are underestimates due to difficulty tracking ligands as they approach the center of the SLB - T cell junction. (**F**) Bound ligand and all ligand under a cell footprint are imaged with long and short exposure times, respectively. Scale bar 5 µm. (**G**) The fraction of Fab’-DNA ligands bound under T cells is very high all Fab’-DNA constructs. Colored bar: median; black square: mean; box: interquartile range; whiskers: data within 1.5x IQR. (H57: n = 27; 2C11: n = 58; 17A2: n = 26). Data are compiled from cells from at least two mice for each condition in dwell time and fraction bound measurements.

NFAT and LAT images were processed in ilastik (55), a machine learning program for bio-image analysis. NFAT-mCherry images were segmented into cytosol and nucleus by training ilastik’s pixel classification algorithm using color/intensity, edge, and texture features. Activated cells were defined as cells with a background-subtracted nucleus intensity to cytosol intensity ratio greater than 1. Only cells well-spread on the bilayer and with clear nuclei were included in analysis. LAT-eGFP images were processed in ilastik to identify LAT condensates. Pixel probability maps corresponding to LAT condensates were imported from ilastik to FIJI to be tracked in TrackMate. The number of LAT condensates experienced by a cell was determined by counting the number of LAT condensate tracks that persisted for at least 4 frames, with a 2 s time lapse between frames.

## RESULTS

### Fab’-DNA constructs are monovalent when conjugated to supported membranes

Fab’-DNA constructs were synthesized by digesting three commercially available anti-murine TCR antibodies: H57-597; 145-2C11; and 17A2, purifying the monovalent Fab’ fragments, and conjugating those fragments to DNA oligonucleotides labeled with a single Atto647N dye, as described previously (see Materials and Methods for details) (Fig. S1 and S2 in the Supporting Material) (25).

Supported lipid bilayers (95% DOPC, 3% MCC-DOPE, and 2% Ni-NTA-DOGS phospholipids) were formed in imaging chambers and functionalized with a thiol-modified DNA oligonucleotide (thiol-DNA) to which the Fab’-DNA molecules could hybridize. SLBs were first formed directly from unmodified SUVs followed by addition of the thiol-DNA, which became covalently linked to the MCC-DOPE lipids. The height of the Fab’-DNA above the bilayer was controlled by the length of this thiol-DNA strand. The shortest thiol-DNA strand was 16 nucleotides and precisely complemented the Fab’ strand. Longer thiol-DNA strands, able to increase the intermembrane space at binding events (56, 57), were created by adding a 19 nt and 59 nt poly(dT) tail to the SLB-proximal 5’ end (Fig 1C). The length of the thiol-DNA strands were designed to allow for up to 14 nm, 25 nm, or 50 nm of vertical space between the SLB and the T cell plasma membrane at binding events between Fab’-DNA and TCR (Fig. S3). Membrane-anchored poly(dT) tails are an established method for generating spacers able to extend past the 20+ nm thick glycocalyx on cell surfaces (56).

The SLB was then functionalized with Fab’-DNA and the adhesion molecule ICAM-1, which binds the integrin receptor LFA-1 and is critical to forming a continuous contact between T cells and antigen presenting cells. Fab’-DNA rapidly anneals to the complementary thiol-DNA (Fig. 2A) and ICAM-1 couples to Ni-NTA lipids through multivalent interactions between Ni^2+^ and its histidine tag (51,65). Though the Atto647N dye used to label Fab’-DNA has a high membrane interaction factor (58), it has previously been confirmed that Fab’-DNA does not interact with the SLB in the absence of the complementary thiol-DNA (25).

H57, 2C11, and 17A2 Fab’-DNA constructs were confirmed to be monovalently conjugated to the SLB using single molecule total internal reflection fluorescence (TIRF) microscopy. For measurement of freely diffusing Fab’-DNA, images were taken with short exposure time (20 ms) and high power (8.6 mW). In streaming image acquisitions (Movies S1 and S2), particles of uniform brightness diffuse across the SLB with uniform two-dimensional Brownian motion (Fig. 2B) and bleach in a single step (Fig. 2C). The FIJI plugin TrackMate was used to link images of single fluorophores into trajectories of diffusing particles, and the step size distribution of the resulting tracks were analyzed to obtain the diffusion coefficient of each Fab’-DNA construct. The step size distributions for both 2C11 Fab’-DNA and 17A2 Fab’-DNA, constructed using a multiple delay time protocol (see Supporting Materials and Methods), were each well-fit by a single species Brownian diffusion model (Figs. 2D, 2F, Fig. S4):

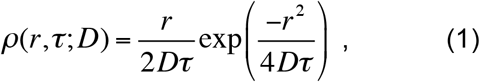

with molecular displacement, *r*, corresponding delay time, *τ*, and diffusion coefficient, D. D values for 2C11 Fab’-DNA and 17A2 Fab’-DNA were measured to be 2.49 ± 0.02 µm^2^/s and 2.83 ± 0.02 µm^2^/s, respectively, in close agreement with the previously reported value of D = 2.43 ± 0.02 µm^2^/s for H57 Fab’-DNA on comparable supported bilayers (25). Step photobleaching analysis further confirmed that all Fab’-DNA ligands conjugated to the SLB as monomers. For both 2C11 and 17A2 Fab’-DNA constructs >99% of fluorescent signals bleached in a single step (Figs. 2E, 2G). The diffusion and step photobleaching data together provide strong evidence that Fab’-DNA exists on the supported membrane as a monovalent, monomeric ligand.

Single particle fluorescence intensity distributions of Fab’-DNA-Atto647N annealed to short (16 nt), medium (36 nt), and long (76 nt) thiol-DNA tethers confirm that the longer tethers create more space between Fab’-DNA and the SLB. In TIRF, illumination intensity decays exponentially from the interface (in this case, between glass and the water layer below the bilayer), and so particle intensity can be a ruler for height for fluorophores if they are in a uniform chemical environment. In this system, the Atto647N dye on Fab’-DNA annealed to the shortest thiol-DNA tether likely intercalates into the membrane, increasing the brightness of the fluorophore and slowing diffusion (25). The intensity distributions of Fab’-DNA-Atto647N annealed to the longer tethers are substantially dimmer, indicating that the Atto647N fluorophore is not able to intercalate into the SLB and that it is at an increased height above the SLB where it experiences a lower illumination intensity (Fig. 2H). As expected, particle intensity decreases as tether length increases. These data agree with the model of the poly(dT) linker as a worm-like chain with a persistence length of 1.5 – 3 nm in 150 mM salt and a fluorophore that can occupy a range of heights above the SLB (59), with the average particle height increasing with linker length. The medium and long thiol-DNA tethers are almost certainly not fully extended when diffusing freely on the bilayer, but they have the capability to extend up to 25 nm and 50 nm, respectively, in response to forces from the apposing T cell membrane.

### All Fab’ ligands bind TCR with high affinity

We then evaluated the interaction between T cells and SLB-tethered agonists by live-cell imaging of primary murine CD4^+^ effector T cells expressing the AND TCR (details in Materials and Methods). T cells were added to imaging chambers containing a continuous SLB decorated with Fab’-DNA and ICAM-1 as described above that was equilibrated on the microscope to 37 °C. T cells made initial contact with the bilayer through interactions of adhesion receptor LFA-1 with ICAM-1, and then spread on the bilayer, creating a junction within which Fab’-DNA molecules could bind TCR (Fig. 3A, Movie S3). This junction was visualized using reflection interference contract microscopy (RICM). When Fab’-DNA binds TCR, its mobility dramatically decreases (Figs. 3B, 3C), as has been reported previously with other TCR ligands (15, 19, 57). Bound ligands are specifically resolved using a long, 500 ms exposure time, which allows slow-moving bound Fab’-DNA to be clearly resolved while blurring the fluorescent signal from quickly diffusing unbound Fab’-DNA (Fig. 3C) (15). Bound Fab’-DNA was tracked through time, with 10 s time lapse between images to minimize photobleaching, enabling direct visualization of individual Fab’-DNA:TCR dwell times (Figs. 3A, 3D), and dwell time distributions for each ligand were assembled from those tracks (Fig. 3E). These distributions were fit with a single exponential decay, based on the first order dissociation kinetics of unbinding. After correcting for photobleaching, all Fab’ ligands exhibited a dwell time, τ_off_, of at least two minutes. In our experiments, Fab’ ligands become difficult to track accurately after about two minutes due to the shuttling of bound TCR to the geometric center of the cell (Fig. 3A), and so these measured dwell times reflect a lower bound. Fab fragments are known to bind strongly to their binding partner, with the H57 Fab fragment reported to bind TCR stably for > 50 min (60). In contrast, the strong agonist MCC-MHC binds AND TCR with a reliably measured dwell time of about 50 s using this method (15, 19, 20, 25).

The Fab’-DNA ligands in this study also bind TCR with a fast on-rate, corresponding to an overall very high efficiency of binding. The total number of Fab’-DNA ligands underneath a cell and the number of TCR-bound Fab’-DNA ligands can be independently measured to determine the fraction of bound Fab’-ligands at any point in time. All Fab’ ligands can be resolved by rapidly acquiring short (50 ms) exposure images while bound Fab’-DNA:TCR complexes can be distinguished using long (500 ms) exposure images, in which the free ligands are diffusing too rapidly to produced well-defined images (Fig. 3F) (19, 25). The cell footprint on the bilayer is determined by RICM, and the fraction of ligands under the cell that are bound, a measure of the efficiency of ligand:receptor binding, can then be directly calculated. The median fraction bound was above 0.85 for all Fab’-DNA constructs, indicating that they all bind AND TCR very efficiently (Fig. 3G). By comparison, about 30% of the strong agonist MCC-MHC, is observed bound to AND TCR at similar ligand densities (19).

We considered the possibility that Fab’-DNA constructs with longer tethers may have slower kinetic on rates due to their increased conformational degrees of freedom and therefore could have a lower measured fraction bound. This, however, was not the case; the measured fraction bound for H57 Fab’-DNA with the medium length (36 nt) and long (76 nt) DNA tethers were almost identical to the fraction of bound H57 Fab’-DNA with the short (16 nt) tether (Fig. S5).

### Fab’-DNA potency is independent of binding epitope, but varies with DNA tether length

We characterized Fab’-DNA potency by measuring T cell activation vs. ligand density dose-response curves. Nuclear localization of the transcription factor nuclear factor for the activation of T cells (NFAT) is a reliable, binary indicator of early T cell activation and has been used as a quantitative readout for the activation of the calcium signaling pathway in previous precision ligand density titrations (19, 20, 25). An NFAT localization reporter, lacking the DNA binding domain and fluorescently tagged with mCherry, allows for facile visualization of NFAT localization without modulating transcriptional activity (42). T cells transduced with the NFAT reporter were added to SLBs presenting ICAM-1 and Fab’-DNA or the strong pMHC agonist, MCC-MHC. Cells were visualized landing and spreading on the bilayer, binding Fab’-DNA, and within minutes, translocating NFAT from the cytosol to the nucleus (Fig. 4A). Cells were defined as activated if the fluorescence intensity from the NFAT reporter was greater in the nucleus than in the cytosol (ratio > 1) (Fig. 4B). The potency of each ligand was measured by counting the fraction of activated cells 20 min after adding cells to bilayers with precisely quantified ligand density (Fig. S6A). Regardless of ligand binding epitope, all ligands that allowed about 14 nm spacing between the SLB and T cell plasma membrane at binding events, MCC-MHC and all Fab’-DNA constructs with 16 nt DNA tether, reached a half-maximal response at a ligand density of 0.2 – 0.4 µm^-2^ (Fig. 4C). Additionally, we have previously shown that short-tethered H57 Fab’-DNA and MCC-MHC stimulate similar IL-2 responses from T cells, indicating that later functional outputs of T cell activation are independent of ligand binding site (25). By contrast, ligand potency was strongly influenced by the length of the thiol-DNA tether. Ligands with DNA tethers that allowed about 25 nm and 50 nm of space between the SLB and T cell plasma membrane were about 10 times and 100 times less potent, respectively, than the ligands that allowed an intermembrane space of about 14 nm (Figs. 4D, S6B). Interestingly, even very high density (∼100 µm^-2^) of the longer tethered ligands did not lead to maximal activation of the T cell population (Fig. S6C).

**Figure 4.**
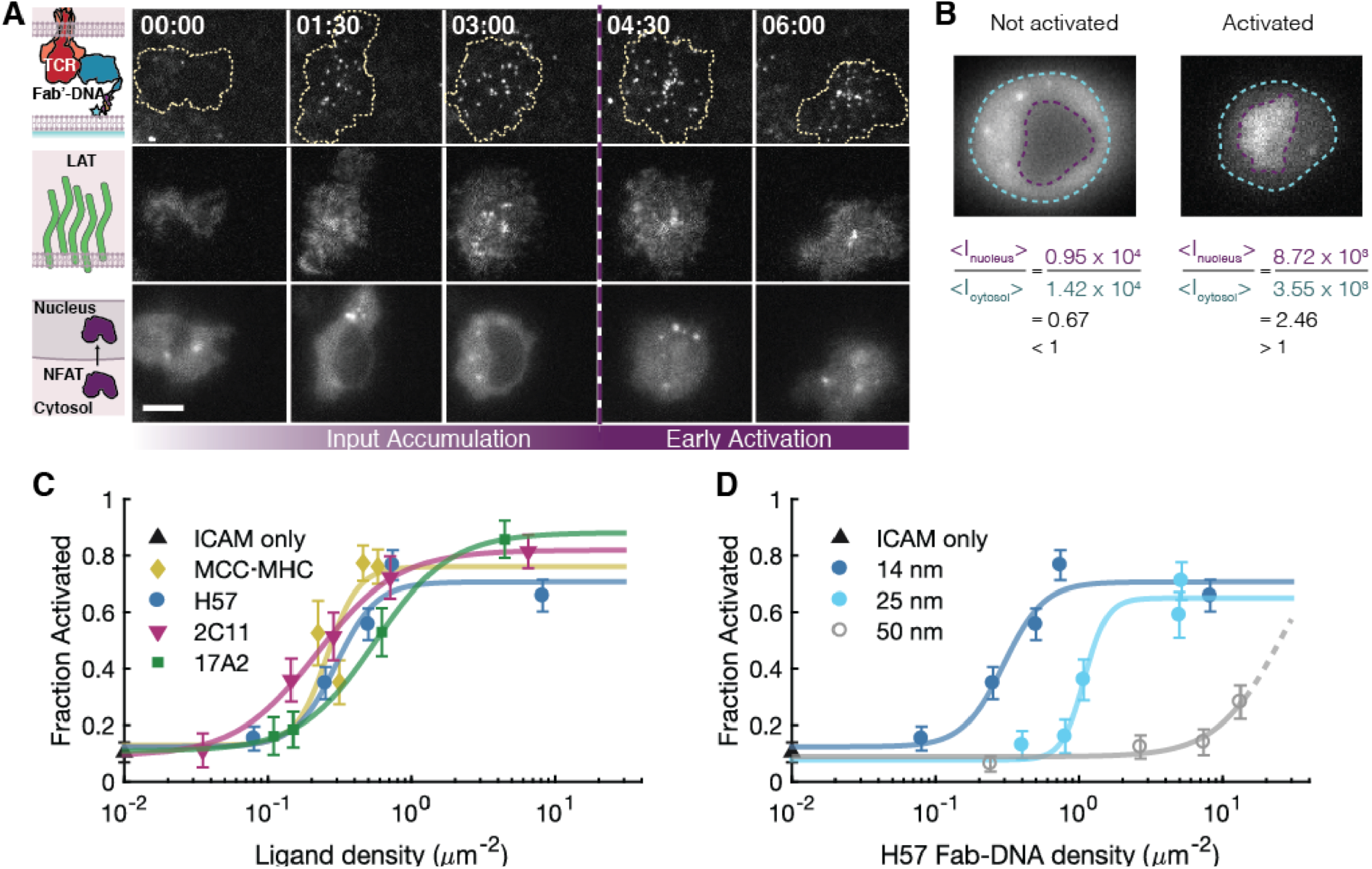
T cells activate with similar probabilities regardless of ligand binding epitope, but have lower activation response when stimulated with Fab’-DNA ligands with longer tethers. (**A**) Cells added to supported lipid bilayers make adhesion contacts, bind TCR ligand (top row), respond to TCR triggering, visualized by LAT condensate formation (middle row), and may activate, visualized by NFAT reporter fluorescence intensity accumulating in the nucleus. Scale bar 5 µm. (**B**) Cells are defined as activated if the ratio of fluorescence intensity in the nucleus to the cytosol is greater than one and not activated if the ratio is less than one. Scale bar 5 µm. (**C** and **D**) Dose-response curves for each ligand are built by adding cells to bilayers with precisely quantified density of ligand, letting cells interact with the bilayers for 20 min, then determining the fraction of activated cells for each condition. n > 50 cells for all conditions. Error bars denote the SEM. Data are representative of at least two biological replicates. (**C**) All ligands that create close, ∼14 nm intermembrane spaces at binding events activate with similar potency, regardless of ligand binding epitope. (**D**) H57 Fab’-DNA potency significantly decreases as the length of the DNA tether increases. The dashed line for longest tether shows the extrapolated fit if a maximum T cell response similar to the shorter ligands is assumed.

### Increased tether length decreases the efficiency of LAT activation by bound TCR

We finally imaged TCR-proximal signaling events to investigate whether the low potencies of long-tethered Fab’-DNA constructs are rooted in a poor ability to trigger TCR efficiently despite robust binding. To do this, we visualized condensation of LAT, which results from phosphorylation activity immediately downstream of triggered TCR (43, 46). LAT condensates have recently been shown to occur even in response to single ligation events between agonist pMHC and TCR (45), and so here, LAT condensation serves as a measure of TCR-proximal signaling activity from individual Fab’-DNA:TCR binding events. T cells expressing LAT-eGFP were introduced to fluid supported lipid bilayers decorated with ICAM-1 and either a Fab’-DNA construct or the agonist MCC-MHC. Using multichannel TIRF microscopy, binding events were imaged as described above, using a long exposure time at low power to specifically image slowly moving ligand, and LAT-eGFP was imaged using low power and moderate exposure time to best capture the dynamic range of LAT fluorescent intensity in the cell being imaged. Cells were imaged starting upon initial contact with the bilayer and continuing for 120 s, during which LAT clustering in response to binding is most active, with a 2 s time lapse between successive frames. The density of Fab’-DNA on bilayers was kept very low (0.01-0.03 µm^-2^) in order to best track single ligation events through time and unambiguously capture the LAT clustering response to these events.

Snapshots from time sequences showed clearly resolved binding events between ligand and TCR and concurrent local increase in LAT density at these binding events for potent ligands (Fig. 5A, first four columns). Less potent ligands – those with longer DNA tethers that allowed for greater intermembrane space – experienced fewer LAT clusters colocalized binding events (Fig. 5A, last two columns). For potent ligands, a single binding event was routinely sufficient to trigger significant LAT clustering (Fig. 5B, circled binding event; Fig. S7A), as has been previously reported for strong pMHC agonists (45). Moreover, T cells form LAT clusters in response to both Fab’-DNA and MCC-MHC when both ligands are presented on the bilayer at low density, indicating that T cells do not readily distinguish ligand identity and that the effects from multiple agonists are additive (Fig. S7B). Interestingly, instances where LAT clusters formed on bilayers presenting weak ligands with long DNA tethers often colocalized with clusters of binding events (Fig. S7C).

**Figure 5.**
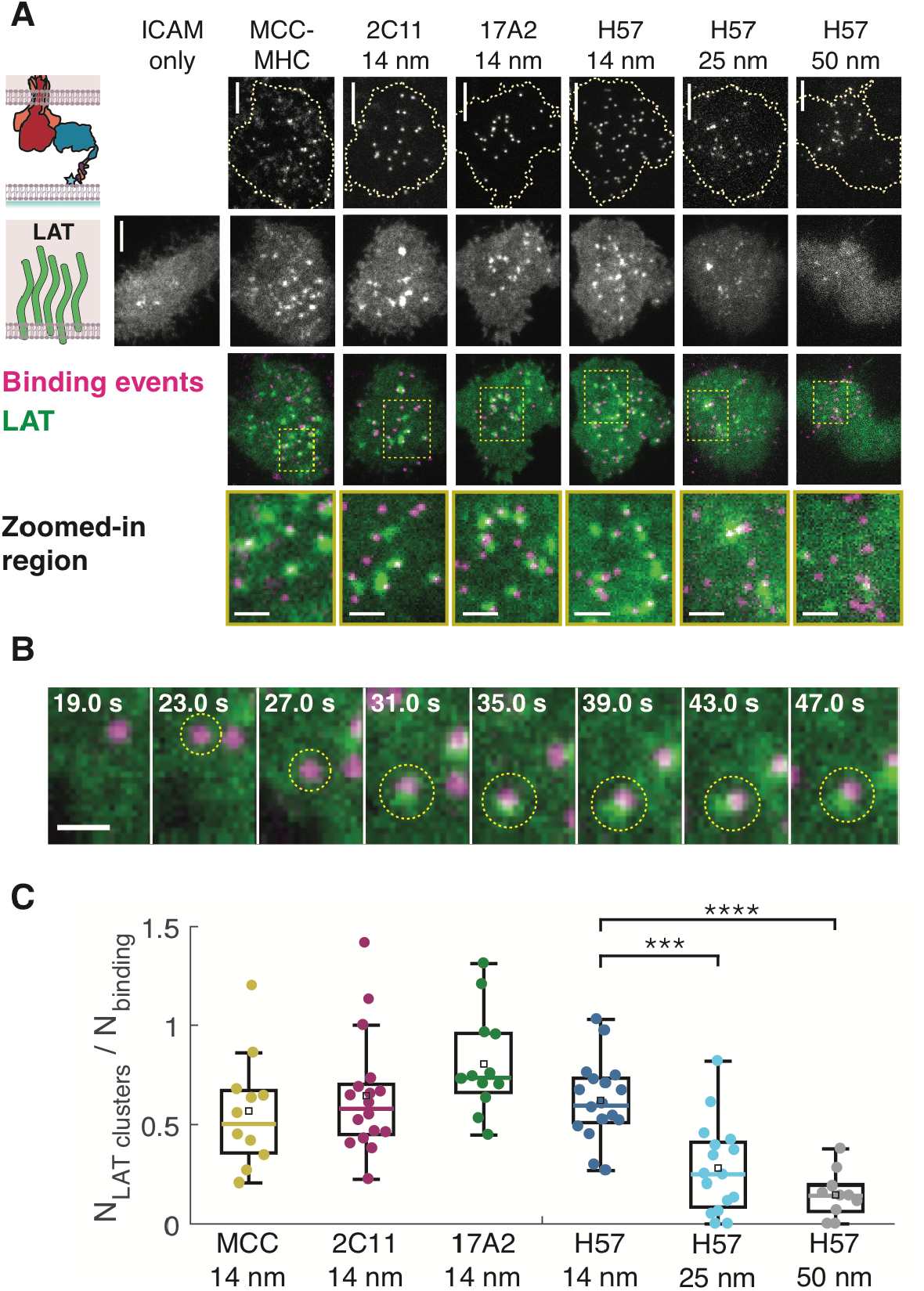
All short ligands trigger TCR with similar efficiency, while ligands with longer tethers trigger TCR with lower efficiency. (**A**) Snapshots of binding events underneath a T cell (top row) and concurrent image of LAT-eGFP (second row) show that LAT condensates often colocalize with binding events from ligands that create a 14 nm intermembrane space, whereas LAT condensates rarely colocalize with binding events from ligands with longer tethers and there are far fewer LAT condensates in these cells (third and fourth rows). Binding events are colored magenta and LAT is colored green. Scale bar for first three rows 5 µm. Scale bar for zoomed-in region in fourth row 2 µm. (**B**) A single binding event between Fab’-DNA and TCR is sufficient to create a proximal LAT condensate. The binding event that appears 23 s after cell landing (yellow circle) triggers the formation of a LAT condensate a handful of seconds later. Scale bar 1 µm. (**C**) The numbers of LAT clusters and binding events experienced by cells were counted for the first 2 min after initial cell contact with the bilayer. There is no significant difference in the ratio of the number of LAT clusters to the number of binding events for short ligands. There is a significant decrease in this ratio for ligands with longer DNA tethers. Colored bar: median; black square: mean; box: interquartile range; whiskers: data within 1.5x IQR. Significance was determined by the Mann-Whitney U-test (*** p < 0.001; **** p < 0.0001; n.s. p > 0.01). Data are compiled from cells from at least two mice for each condition.

To quantify the signaling efficiency of each ligand, the total number of binding events experienced by a cell and the total number of LAT clusters formed within the cell were counted for first two minutes of the cell interacting with the supported membrane. Binding event trajectories were counted after tracking all events in TrackMate (54). LAT clusters were first identified in each frame using ilastik (55), a user-friendly machine learning program for bio-image analysis, and then LAT pixel probability maps produced by ilastik were tracked in TrackMate (details in Supporting Materials and Methods). The ratio of the number of LAT clusters to the number of binding events was then calculated for each cell (Fig. 5C). Some LAT clusters formed in the absence of binding (see ICAM only example in Fig. 5A), and so these ratios occasionally exceed one. In agreement with the NFAT titrations, all ligands that allowed about 14 nm spacing between the SLB and T cell plasma membrane produced LAT clusters with a similar efficiency per binding events, with medians of around 0.6. The ratio of LAT clusters to binding events significantly decreased with increasing intermembrane space (Fig. 5C, Tables S1-S2). These results indicate that the difference in ligand potency seen at the level of transcription factor localization results from differences in the ligands’ abilities to trigger TCR.

## DISCUSSION

### Implications for further Fab’-DNA development

Fab’-DNA shows promise as a tool to activate of polyclonal T cell populations in a manner similar to agonist pMHC; Fab’-DNA is monovalent and activates T cells at low ligand densities when membrane-associated but is inactive from solution (25). Here, we use the modular design of Fab’-DNA to test the dependence of T cell response on Fab’-DNA binding epitope and tether length. To achieve the goal of designing a universal TCR agonist, Fabs used in Fab’-DNA constructs must bind the constant region of the TCR, which is far from the pMHC binding site at the apex of the TCR variable region (Fig. 1B). The robust result that signaling from the TCR is indifferent to binding site allows flexibility in Fab’ choice when using anti-murine Fab’-DNA and in designing Fab’-DNA constructs capable of binding and activating human T cells or perhaps chimeric antigen receptor T cells. That signaling potency depends on the spacing between apposed membranes at binding events is consistent with previous work with pMHC and therapeutic TCR-binding molecules (38, 47, 48).

The experimental system described herein is similar to a recent study by Roffler and coworkers that found that very high affinity (< 1 nM) anti-CD3 single chain variable fragments (scFv) did not exhibit a tether-length dependence on T cell activation (61). However, an anti-CD3 scFv with 8 nM affinity exhibited an order of magnitude decrease in potency as the intermembrane spacing at binding events increased from 10 to 40 nm, and 2C11 scFv, with a reported affinity of 70 nM, exhibited a dramatic decrease in potency as tether length increased, in close agreement with the results presented in our study. Notably, moderate-affinity antibody fragments bind TCR/CD3 considerably more strongly than pMHC, and further work developing Fab’-DNA to mimic pMHC would focus on further decreasing, not increasing, Fab’ binding affinity, potentially enhancing the tether-length dependence of these ligands. The density-dependent data presented in Chen *et al*., reporting that the T cell response is nearly identical when interacting with APCs decorated with 10 µm^-2^ compared to 100 µm^-2^ of their shortest ligand (61), also agree with our data, which show that the threshold activation density for short ligands is around 0.2 µm^-2^, near physiological pMHC densities, and T cell populations are fully activated by about 1 µm^-2^ ligand.

### Relating results to proposed mechanisms of T-cell receptor triggering

The data presented in this study most closely relate to two proposed (and by no means mutually exclusive) mechanisms of TCR triggering: (1) kinetic-segregation and (2) anisotropic mechanosensing by the TCR. The kinetic-segregation model proposes that when TCR is localized in close contact zones between apposed membranes for a sufficient length of time, whether by ligand binding or not, the bulky and abundant phosphatase CD45 is excluded from the proximity of TCR (28–32, 62, 63). CD45 exclusion increases the probability that TCR and proximal downstream kinases and scaffolding molecules are phosphorylated and leads to signal trandsuction (64). All results presented herein are consistent with kinetic-segregation. Interpreting the data according to this model, short Fab’-DNA ligands are able to pin TCR in close contact zones that exclude CD45 for extended lengths of time, even if that zone is created by only a single binding event. Increasing the DNA tether length prevents the formation of close contact zones at binding events and increases the availability of bound TCR to CD45.

The relation of these data to the mechanosensing model is more complex. The model of the TCR as an anisotropic mechanosensor proposes that ligands able to exert a certain torque on the TCR are more effective at triggering TCR than ligands incapable of applying such torque. According to this model, force from pMHC binding is transmitted through the FG loop of TCRβ, causes allosteric change in the TCR/CD3 constant regions, and brings TCR/CD3 to a signaling-competent state (27, 65). H57 binds the FG loop of TCRβ, and is thought to trigger TCR by directly acting on this hinge when its Fab fragment is bound to a surface (34). 2C11 has two potential binding sites on the TCR/CD3 complex, as each complex has two CD3ε subunits. One CD3ε subunit sits directly above the FG loop, and 2C11 binding to this subunit has been proposed to act on the FG loop hinge similarly to H57 (26). The second CD3ε subunit sits approximately 120° away from the first (66). The observation that 2C11 Fab’-DNA has a similar potency as H57 Fab’-DNA and MCC-MHC suggests that 2C11 Fab’-DNA is able to trigger TCR regardless of which CD3ε subunit it binds. This interpretation is corroborated by 2C11 Fab’-DNA:TCR binding events producing LAT clusters with the same efficiency as H57 Fab’-DNA and MCC-MHC. Due to the relatively low densities of Fab’-DNA used in these studies (0.01 - 1 µm^-2^), it is unlikely that two 2C11 Fab’-DNA molecules bind the same TCR.

The results from 17A2 Fab’-DNA, matching results from all ligands that bind other epitopes, further underscore that TCR/CD3 appears to be indifferent to where it is bound. 17A2 binds the CD3ε/CD3γ cleft, off-axis from the FG loop fulcrum (26, 66). In experiments by Kim and coworkers, using optical tweezers and in the absence of adhesion, 17A2 antibody activated T cells only with the exertion of a tangential force. However, the experimental data we present here are from supported lipid bilayer assays with low 17A2 Fab’-DNA ligand density and in the context of physiologically dense ICAM-1:LFA-1 adhesion. In this configuration, Fab’-DNA:TCR binding always occurs within a few tens of nanometers of ICAM-1:LFA-1 adhesion complexes, and is not mechanically isolated from these (67, 68). Mechanical coupling among protein complexes within the intermembrane junction may obviate the previously observed requirement of directional torque to trigger TCR. Notably, free lateral diffusion of ligands in the SLB prevents sustained tangential forces, and normal force applied to a single half antibody at a T cell–bead interface has been shown to be suffienct to activate T cells (69). It does not appear that TCR/CD3 is sensitive to which region or regions of the complex bear the load of a normal force in context of ICAM-1:LFA-1 adhesion.

These experiments, however, do not rule out force as an important mediator of TCR triggering by Fab’-DNA. TCRs bound the Fab’-DNA with short DNA tethers are more likely to sustain normal forces, due to the presence of larger ICAM-1:LFA-1 binding interactions and proteins on the T cell with large extracellular domains like CD45, compared to TCRs bound by Fab’-DNA with longer tethers. It is possible that the forces on TCR bound to short ligand are greater in our experimental platform compared to a junction between an antigen presenting cell and a T cell because small adhesion proteins such as CD2 are not included in our bilayers.

### Binding of a single Fab’-DNA is capable of creating significant TCR-proximal signaling

Our experiments are designed for facile, direct observation of single binding events between ligand and TCR with the ability to simultaneously image signaling response. With this design, we routinely see LAT condensates triggered by single ligand:TCR binding events with both pMHC (45) and Fab’-DNA ligands. These data stand in apparent contrast to a recently published, and carefully executed, study by Sevscik and coworkers that concludes that a similar monovalent H57-derived ligand, reported to allow 12-19 nm space between the SLB and T cell, activates T cells as efficiently as MCC-MHC only if two ligands are within a lateral distance of 20 nm from each other (70). Interestingly, in our study, the weaker Fab’-DNA ligands with longer DNA tethers were measured to be more likely to generate a LAT condensate if multiple ligands are bound in a diffraction-limited area. This observation, that less potent ligands cause TCR-proximal signaling more readily when they are clustered, may relate to other studies that report ligand clustering as a requirement for signaling using similar experimental platforms (57, 70).

In conclusion, through modulating the binding epitope and DNA tether length of Fab’-DNA constructs, we find that varying the DNA tether length affects Fab’-DNA potency, but varying the binding epitope does not. These findings advance the utility of membrane-linked Fab’-DNA ligands as universal T cell agonists and clarify design criteria for further development of this class of synthetic T call activators.

## Acknowledgements

We are grateful to members of the Groves Laboratory for critical feedback on this manuscript. We thank F. Marangoni (Harvard Medical School) for providing the NFAT reporter plasmid. We thank L. Teyton (Scripps Research Institute) and M. Davis (Stanford University) for providing the MHC and ICAM-1 bacmids. We thank Scott Hansen (University of Oregon) for cloning the NFATmCherry-P2A-LAT-eGFP construct.

The financial support for this work was provided by National Institutes of Health grant P01 AI091580 and by the Novo Nordisk Foundation Challenge Programme as part of the Center for Geometrically Engineered Cellular Systems.

## Author Contributions

JTG and KBW conceived and designed this study. KBW and SM performed experiments. KBW, SM and DMB contributed new reagents/analytical tools. SK, MKO’D, KBW, DMB, and SM harvested and prepared T cells for experiments. KBW analyzed data. KBW and JTG wrote the manuscript. All authors edited the manuscript.

## REFERENCES

1. Weiss, A., J. Imboden, K. Hardy, B. Manger, C. Terhorst, and J. Stobo. 1986. The Role of the T3/Antigen Receptor Complex in T-Cell Activation. Annu. Rev. Immunol. 4:593–619.

2. Kaye, J., S. Porcelli, J. Tite, B. Jones, and C.A. Janeway. 1983. Both a monoclonal antibody and antisera specific for determinants unique to individual cloned helper T cell lines can substitute for antigen and antigen-presenting cells in the activation of T cells. J. Exp. Med. 158:836–856.

3. Kubo, R.T., W. Born, J.W. Kappler, P. Marrack, and M. Pigeon. 1989. Characterization of a monoclonal antibody which detects all murine alpha beta T cell receptors. J. Immunol. 142:2736–42.

4. Leo, O., M. Foo, D.H. Sachs, L.E. Samelson, and J.A. Bluestone. 1987. Identification of a monoclonal antibody specific for a murine T3 polypeptide. Proc. Natl. Acad. Sci. U. S. A. 84:1374–1378.

5. Miescher, G.C., M. Schreyer, and H.R. MacDonald. 1989. Production and characterization of a rat monoclonal antibody against the murine CD3 molecular complex. Immunol. Lett. 23:113–118.

6. Irving, B.A., and A. Weiss. 1991. The cytoplasmic domain of the T cell receptor ζ chain is sufficient to couple to receptor-associated signal transduction pathways. Cell. 64:891–901.

7. Boniface, J.J., J.D. Rabinowtiz, C. Wulfing, J. Hampi, Z. Reich, J.D. Altman, R.M. Kantor, C. Beeson, H.M. McConnell, and M.M. Davis. 1998. Initiation of signal transduction through the T cell receptor requires the multivalent engagement of peptide/MHC ligands. Immunity. 9:459–466.

8. Kaye, J., and C.A. Janeway. 1984. The Fab fragment of a directly activating monoclonal antibody that precipitates a disulfide-linked heterodimer from a helper T cell clone blocks activation by either allogeneic Ia or antigen and self-Ia. J. Exp. Med. 159:1397–1412.

9. Altman, J.D., P.A.H. Moss, P.J.R. Goulder, D.H. Barouch, M.G. McHeyzer-Williams, J.I. Bell, A.J. McMichael, and M.M. Davis. 1996. Phenotypic analysis of antigen-specific T lymphocytes. Science. 274:94–96.

10. Johnson, K.G., S.K. Bromley, M.L. Dustin, and M.L. Thomas. 2000. A supramolecular basis for CD45 tyrosine phosphatase regulation in sustained T cell activation. Proc. Natl. Acad. Sci. 97:10138–10143.

11. Ledbetter, J.A., C.H. June, L.S. Grosmaire, and P.S. Rabinovitch. 1987. Crosslinking of surface antigens causes mobilization of intracellular ionized calcium in T lymphocytes. Proc. Natl. Acad. Sci. 84:1384–1388.

12. Irvine, D.J., M.A. Purbhoo, M. Krogsgaard, and M.M. Davis. 2002. Direct observation of ligand recognition by T cells. Nature. 419:845–849.

13. Purbhoo, M.A., D.J. Irvine, J.B. Huppa, and M.M. Davis. 2004. T cell killing does not require the formation of a stable mature immunological synapse. Nat. Immunol. 5:524–530.

14. Huang, J., M. Brameshuber, X. Zeng, J. Xie, Q. jing Li, Y. hsiu Chien, S. Valitutti, and M.M. Davis. 2013. A Single peptide-major histocompatibility complex ligand triggers digital cytokine secretion in CD4+ T Cells. Immunity. 39:846–857.

15. O’Donoghue, G.P., R.M. Pielak, A.A. Smoligovets, J.J. Lin, and J.T. Groves. 2013. Direct single molecule measurement of TCR triggering by agonist pMHC in living primary T cells. Elife. 2:e00778.

16. Demotz, S., H.M. Grey, and A. Sette. 1990. The Minimal Number of Class II MHC-Antigen Complexes Needed for T Cell Activation. Science. 249:1028–1030.

17. Harding, C. V., and E.R. Unanue. 1990. Quantitation of antigen-presenting cell MHC class II/peptide complexes necessary for T-cell stimulation. Nature. 346:574–576.

18. Bozzacco, L., H. Yu, H.A. Zebroski, J. Dengjel, H. Deng, S. Mojsov, and R.M. Steinman. 2011. Mass spectrometry analysis and quantitation of peptides presented on the MHC II molecules of mouse spleen dendritic cells. J. Proteome Res. 10:5016–5030.

19. Pielak, R.M., G.P. O’Donoghue, J.J. Lin, K.N. Alfieri, N.C. Fay, S.T. Low-Nam, and J.T. Groves. 2017. Early T cell receptor signals globally modulate ligand: receptor affinities during antigen discrimination. Proc. Natl. Acad. Sci. U. S. A. 114:12190–12195.

20. Lin, J.J.Y., S.T. Low-Nam, K.N. Alfieri, D.B. McAffee, N.C. Fay, and J.T. Groves. 2019. Mapping the stochastic sequence of individual ligand-receptor binding events to cellular activation: T cells act on the rare events. Sci. Signal. 12:1–14.

21. Varma, R., G. Campi, T. Yokosuka, T. Saito, and M.L. Dustin. 2006. T Cell Receptor-Proximal Signals Are Sustained in Peripheral Microclusters and Terminated in the Central Supramolecular Activation Cluster. Immunity. 25:117–127.

22. Mossman, K.D., G. Campi, J.T. Groves, and M.L. Dustin. 2005. Altered TCR Signaling from Geometrically Repatterned Immunological Synapses. Science. 310:1191–1193.

23. Monks, C.R.F., B.A. Freiberg, H. Kupfer, N. Sciaky, and A. Kupfer. 1998. Three-dimensional segregation of supramolecular activation clusters in T cells. Nature. 395:82–86.

24. Grakoui, A., S.K. Bromley, C. Sumen, M.M. Davis, A.S. Shaw, P.M. Allen, and M.L. Dustin. 1999. The immunological synapse: A molecular machine controlling T cell activation. Science. 285:221–227.

25. Lin, J.J., G.P. O’Donoghue, K.B. Wilhelm, M.P. Coyle, S.T. Low-Nam, N.C. Fay, K.N. Alfieri, and J.T. Groves. 2020. Membrane Association Transforms an Inert Anti-TCRβ Fab’ Ligand into a Potent T Cell Receptor Agonist. Biophys. J. 118:2879–2893.

26. Kim, S.T., K. Takeuchi, Z.Y.J. Sun, M. Touma, C.E. Castro, A. Fahmy, M.J. Lang, G. Wagner, and E.L. Reinherz. 2009. The αβ T cell receptor is an anisotropic mechanosensor. J. Biol. Chem. 284:31028–31037.

27. Brazin, K.N., R.J. Mallis, D.K. Das, Y. Feng, W. Hwang, J. huai Wang, G. Wagner, M.J. Lang, and E.L. Reinherz. 2015. Structural features of the aβTCR mechanotransduction apparatus that promote pMHC discrimination. Front. Immunol. 6:441.

28. Davis, S.J., and P.A. van der Merwe. 1996. The structure and ligand interactions of CD2: implications for T-cell function. Immunol. Today. 17:177–187.

29. Davis, S.J., and P.A. van der Merwe. 2006. The kinetic-segregation model: TCR triggering and beyond. Nat. Immunol. 7:803–809.

30. Dushek, O., J. Goyette, and P.A. van der Merwe. 2012. Non-catalytic tyrosine-phosphorylated receptors. Immunol. Rev. 250:258–276.

31. James, J.R., and R.D. Vale. 2012. Biophysical mechanism of T-cell receptor triggering in a reconstituted system. Nature. 487:64–69.

32. Chang, V.T., R.A. Fernandes, K.A. Ganzinger, S.F. Lee, C. Siebold, J. McColl, P. Jönsson, M. Palayret, K. Harlos, C.H. Coles, E.Y. Jones, Y. Lui, E. Huang, R.J.C. Gilbert, D. Klenerman, A.R. Aricescu, and S.J. Davis. 2016. Initiation of T cell signaling by CD45 segregation at “close contacts.” Nat. Immunol. 17:574–582.

33. Wang, J.H., K. Lim, A. Smolyar, M.K. Teng, J.H. Liu, A.G.D. Tse, J. Liu, R.E. Hussey, Y. Chishti, C.T. Thomson, R.M. Sweet, S.G. Nathenson, H.C. Chang, J.C. Sacchettini, and E.L. Reinherz. 1998. Atomic structure of an αβ T cell receptor (TCR) heterodimer in complex with an anti-TCR Fab fragment derived from a mitogenic antibody. EMBO J. 17:10–26.

34. Das, D.K., Y. Feng, R.J. Mallis, X. Li, D.B. Keskin, R.E. Hussey, S.K. Brady, J.H. Wang, G. Wagner, E.L. Reinherz, and M.J. Lang. 2015. Force-dependent transition in the T-cell receptor β-subunit allosterically regulates peptide discrimination and pMHC bond lifetime. Proc. Natl. Acad. Sci. U. S. A. 112:1517–1522.

35. Hwang, W., R.J. Mallis, M.J. Lang, and E.L. Reinherz. 2020. The αβTCR mechanosensor exploits dynamic ectodomain allostery to optimize its ligand recognition site. Proc. Natl. Acad. Sci. U. S. A. 117:21336–21345.

36. Garboczi, D.N., P. Ghosh, U. Utz, Q.R. Fan, W.E. Biddison, and D.C. Wiley. 1996. Structure of the complex between human T-cell receptor, viral peptide, and HLA-A2. Nature. 384:134–141.

37. Garcia, K.C., M. Degano, L.R. Pease, M. Huang, P.A. Peterson, L. Teyton, and I.A. Wilson. 1998. Structural Basis of Plasticity in T Cell Receptor Recognition of a Self Peptide-MHC Antigen. Science. 279:1166–1172.

38. Choudhuri, K., D. Wiseman, M.H. Brown, K. Gould, and P.A. Van Der Merwe. 2005. T-cell receptor triggering is critically dependent on the dimensions of its peptide-MHC ligand. Nature. 436:578–582.

39. Springer, T.A. 1990. Adhesion Receptors of the Immune System. Nature. 346:425–434.

40. Woollett, G.R., A.F. Williams, and D.M. Shotton. 1985. Visualisation by low-angle shadowing of the leucocyte-common antigen. A major cell surface glycoprotein of lymphocytes. EMBO J. 4:2827–2830.

41. McCall, M.N., D.M. Shotton, and A.N. Barclay. 1992. Expression of soluble isoforms of rat CD45. Analysis by electron microscopy and use in epitope mapping of anti-CD45R monoclonal antibodies. Immunology. 76:310–7.

42. Marangoni, F., T.T. Murooka, T. Manzo, E.Y. Kim, E. Carrizosa, N.M. Elpek, and T.R. Mempel. 2013. The Transcription Factor NFAT Exhibits Signal Memory during Serial T Cell Interactions with Antigen-Presenting Cells. Immunity. 38:237–249.

43. Zhang, W., J. Sloan-Lancaster, J. Kitchen, R.P. Trible, and L.E. Samelson. 1998. LAT: The ZAP-70 tyrosine kinase substrate that links T cell receptor to cellular activation. Cell. 92:83–92.

44. Nag, A., M.I. Monine, J.R. Faeder, and B. Goldstein. 2009. Aggregation of Membrane Proteins by Cytosolic Cross-Linkers: Theory and Simulation of the LAT-Grb2-SOS1 System. Biophys. J. 96:2604–2623.

45. Ganti, R.S., W.-L. Lo, D.B. McAffee, J.T. Groves, A. Weiss, and A.K. Chakraborty. 2020. How the T cell signaling network processes information to discriminate between self and agonist ligands. Proc. Natl. Acad. Sci. 117:26020–26030.

46. Balagopalan, L., R.L. Kortum, N.P. Coussens, V.A. Barr, and L.E. Samelson. 2015. The Linker for Activation of T Cells (LAT) Signaling Hub: From Signaling Complexes to Microclusters. J. Biol. Chem. 290:26422–26429.

47. Bluemel, C., S. Hausmann, P. Fluhr, M. Sriskandarajah, W.B. Stallcup, P.A. Baeuerle, and P. Kufer. 2010. Epitope distance to the target cell membrane and antigen size determine the potency of T cell-mediated lysis by BiTE antibodies specific for a large melanoma surface antigen. Cancer Immunol. Immunother. 59:1197–1209.

48. Li, J., N.J. Stagg, J. Johnston, M.J. Harris, S.A. Menzies, D. DiCara, V. Clark, M. Hristopoulos, R. Cook, D. Slaga, R. Nakamura, L. McCarty, S. Sukumaran, E. Luis, Z. Ye, T.D. Wu, T. Sumiyoshi, D. Danilenko, G.Y. Lee, K. Totpal, D. Ellerman, I. Hötzel, J.R. James, and T.T. Junttila. 2017. Membrane-Proximal Epitope Facilitates Efficient T Cell Synapse Formation by Anti-FcRH5/CD3 and Is a Requirement for Myeloma Cell Killing. Cancer Cell. 31:383–395.

49. Rangarajan, S., Y. He, Y. Chen, M.C. Kerzic, B. Ma, R. Gowthaman, B.G. Pierce, R. Nussinov, R.A. Mariuzza, and J. Orban. 2018. Peptide–MHC (pMHC) binding to a human antiviral T cell receptor induces long-range allosteric communication between pMHC- and CD3-binding sites. J. Biol. Chem. 293:15991–16005.

50. Coyle, M.P., Q. Xu, S. Chiang, M.B. Francis, and J.T. Groves. 2013. DNA-mediated assembly of protein heterodimers on membrane surfaces. J. Am. Chem. Soc. 135:5012–5016.

51. Kaye, J., M.-L. Hsu, M.-E. Sauron, S.C. Jameson, N.R.J. Gascoigne, and S.M. Hedrick. 1989. Selective development of CD4+ T cells in transgenic mice expressing a class II MHC-restricted antigen receptor. Nature. 341:746–749.

52. Smith, A.W., A.A. Smoligovets, and J.T. Groves. 2011. Patterned Two-Photon Photoactivation Illuminates Spatial Reorganization in Live Cells. J. Phys. Chem. A. 115:3867–3875.

53. Edelstein, A., N. Amodaj, K. Hoover, R. Vale, and N. Stuurman. 2010. Computer control of microscopes using manager. Curr. Protoc. Mol. Biol. 92:14.20.1-14.20.17.

54. Tinevez, J.Y., N. Perry, J. Schindelin, G.M. Hoopes, G.D. Reynolds, E. Laplantine, S.Y. Bednarek, S.L. Shorte, and K.W. Eliceiri. 2017. TrackMate: An open and extensible platform for single-particle tracking. Methods. 115:80–90.

55. Berg, S., D. Kutra, T. Kroeger, C.N. Straehle, B.X. Kausler, C. Haubold, M. Schiegg, J. Ales, T. Beier, M. Rudy, K. Eren, J.I. Cervantes, B. Xu, F. Beuttenmueller, A. Wolny, C. Zhang, U. Koethe, F.A. Hamprecht, and A. Kreshuk. 2019. Ilastik: Interactive Machine Learning for (Bio)Image Analysis. Nat. Methods. 16:1226–1232.

56. Selden, N.S., M.E. Todhunter, N.Y. Jee, J.S. Liu, K.E. Broaders, and Z.J. Gartner. 2012. Chemically programmed cell adhesion with membrane-anchored oligonucleotides. J. Am. Chem. Soc. 134:765–768.

57. Taylor, M.J., K. Husain, Z.J. Gartner, S. Mayor, and R.D. Vale. 2017. A DNA-Based T Cell Receptor Reveals a Role for Receptor Clustering in Ligand Discrimination. Cell. 169:108–119.

58. Hughes, L.D., R.J. Rawle, and S.G. Boxer. 2014. Choose Your Label Wisely: Water-Soluble Fluorophores Often Interact with Lipid Bilayers. PLoS One. 9:e87649.

59. Murphy, M.C., I. Rasnik, W. Cheng, T.M. Lohman, and T. Ha. 2004. Probing Single-Stranded DNA Conformational Flexibility Using Fluorescence Spectroscopy. Biophys. J. 86:2530–2537.

60. Huppa, J.B., M. Axmann, M.A. Mörtelmaier, B.F. Lillemeier, E.W. Newell, M. Brameshuber, L.O. Klein, G.J. Schütz, and M.M. Davis. 2010. TCR–peptide–MHC interactions in situ show accelerated kinetics and increased affinity. Nature. 463:963–967.

61. Chen, B.M., M.A. Al-Aghbar, C.H. Lee, T.C. Chang, Y.C. Su, Y.C. Li, S.E. Chang, C.C. Chen, T.H. Chung, Y.C. Liao, C.H. Lee, and S.R. Roffler. 2017. The affinity of elongated membrane-tethered ligands determines potency of T cell receptor triggering. Front. Immunol. 8:1–21.

62. Carbone, C.B., N. Kern, R.A. Fernandes, E. Hui, X. Su, K.C. Garcia, and R.D. Vale. 2017. In vitro reconstitution of T cell receptor-mediated segregation of the CD45 phosphatase. Proc. Natl. Acad. Sci. U. S. A. 114:E9338–E9345.

63. Fernandes, R.A., K.A. Ganzinger, J.C. Tzou, P. Jönsson, S.F. Lee, M. Palayret, A.M. Santos, A.R. Carr, A. Ponjavic, V.T. Chang, C. Macleod, B. Christoffer Lagerholm, A.E. Lindsay, O. Dushek, A. Tilevik, S.J. Davis, and D. Klenerman. 2019. A cell topography-based mechanism for ligand discrimination by the T cell receptor. Proc. Natl. Acad. Sci. U. S. A. 116:14002–14010.

64. Hui, E., and R.D. Vale. 2014. In vitro membrane reconstitution of the T-cell receptor proximal signaling network. Nat. Struct. Mol. Biol. 21:133–142.

65. Reinherz, E.L. 2019. The structure of a T-cell mechanosensor. Nature. 7–9.

66. Dong, D., L. Zheng, J. Lin, B. Zhang, Y. Zhu, N. Li, S. Xie, Y. Wang, N. Gao, and Z. Huang. 2019. Structural basis of assembly of the human T cell receptor–CD3 complex. Nature. 573:546–552.

67. Qi, S.Y., J.T. Groves, and A.K. Chakraborty. 2001. Synaptic pattern formation during cellular recognition. Proc. Natl. Acad. Sci. 98:6548–6553.

68. Kaizuka, Y., and J.T. Groves. 2006. Hydrodynamic Damping of Membrane Thermal Fluctuations near Surfaces Imaged by Fluorescence Interference Microscopy. Phys. Rev. Lett. 96:118101.

69. Feng, Y., K.N. Brazin, E. Kobayashi, R.J. Mallis, E.L. Reinherz, and M.J. Lang. 2017. Mechanosensing drives acuity of αβ T-cell recognition. Proc. Natl. Acad. Sci. U. S. A. 114:E8204–E8213.

70. Hellmeier, J., R. Platzer, A.S. Eklund, T. Schlichthaerle, A. Karner, V. Motsch, M.C. Schneider, E. Kurz, V. Bamieh, M. Brameshuber, J. Preiner, R. Jungmann, H. Stockinger, G.J. Schütz, J.B. Huppa, and E. Sevcsik. 2021. DNA origami demonstrate the unique stimulatory power of single pMHCs as T cell antigens. Proc. Natl. Acad. Sci. U. S. A. 118:e2016857118.

## SUPPORTING REFERENCES

71. Rousseaux, J., R. Rousseaux-Prévost, H. Bazin, and G. Biserte. 1983. Proteolysis of rat IgG subclasses by Staphylococcus aureus V8 proteinase. Biochim. Biophys. Acta - Protein Struct. Mol. Enzymol. 748:205–212.

72. Hartman, N.C., J.A. Nye, and J.T. Groves. 2009. Cluster size regulates protein sorting in the immunological synapse. Proc. Natl. Acad. Sci. U. S. A. 106:12729–12734.

73. Nye, J.A., and J.T. Groves. 2008. Kinetic control of histidine-tagged protein surface density on supported lipid bilayers. Langmuir. 24:4145–4149.

