## Supporting Materials for "Height, but not binding epitope, affects the potency of synthetic TCR agonists"

**Contents:**

Supporting Materials and Methods

Supporting Figures 1-7

Supporting Tables 1 and 2

Captions for Supporting Movies 1-3

Supporting Materials and Methods

***Reagents***

The phospholipids 1,2-dioleoyl-sn-glycero-3-phosphocholine (DOPC), 1,2-dioleoyl-sn-glycero-3-[(N-(5-amino-1-carboxypentyl)iminodiacetic acid)succinyl] nickel salt (Ni-NTA DOGS), 1,2-dioleoyl-sn-glycero-3-phosphoethanolamine-N-[4-(p-maleimidomethyl)cyclohexane-carboxamide] sodium salt (MCC-DOPE), and 1,2-dipalmitoyl-sn-glycero-3-phosphocholine (DPPC) for preparation of supported lipid bilayer were purchased from Avanti polar lipids (Alabaster, AL, USA) as chloroform solutions. Pepsin from porcine gastric mucosa (3200-4500 U/mg, lyophilized powder) and endoproteinase Glu-C from *Staphylococcus aureus* strain V8 was purchased from Sigma-Aldrich (St. Louis, MO, USA). ATTO647N maleimide was purchased from ATTO-TEC (Irvine, CA, USA), and custom DNA oligonucleotides were purchased from Integrated DNA Technologies (Coralville, IA, USA). Succinimidyl-[(N-maleimidopropionamido)-hexaethyleneglycol] ester (SM-$PEG_6$-maleimide), Protein A, Protein A/G, and Protein G resins were purchased from Thermo Fisher (Waltham, MA, USA). The rat anti-mouse CD3 IgG2b 17A2 clone (InVivoMAb anti-mouse CD3) was purchased from Bio X Cell (Lebanon, NH, USA). The Armenian hamster anti-mouse CD3ε IgG 145-2C11 clone (2C11) was purchased from Biolegend (San Diego, CA). The Armenian hamster anti-mouse TCRβ IgG H57-597 clone (H57) was purchased from Biolegend (San Diego, CA). Sephadex-packed desalting column NAP-5 and PD-10, products from Cytiva (Marlborough, MA, USA), were purchased from standard suppliers. Acrylamide-packed purification columns Bio-Spin 6 and Micro Bio-Spin 6 were purchased from BIO-RAD (Hercules, CA, USA). Centrifugal filters for ultrafiltration Amicon Ultra-0.5, -4, or -15 for several MWCO were purchased from Millipore Sigma (Burlington, MA). RVC medium for T cell culture was prepared with the following composition: DMEM (Gibco, Thermo Fisher) with 10% FBS, 1 mM sodium pyruvate, 2 mM L-glutamine, 1x Corning nonessential amino acids (Fisher Scientific, Thermo Fisher), 1x Corning MEM vitamin solution (Fisher Scientific), 0.67 mM L-arginine, 0.27 mM L-asparagine, 14 uM folic acid, 1x Corning Penicilline/Streptomycin (100 IU, 0.1 mg/mL respectively) (Fisher Scientfic), 50 uM $\beta$-mercaptoethanol. Other common chemical reagents were purchased from standard suppliers.

***Oligonucleotide Sequences***

The sequences of the oligonucleotides used in this study are summarized below. The sequence 1 is the ssDNA conjugated to Fab', and the sequences 2-4 are the ssDNA covalently linked to supported lipid bilayer.

Sequence 1: 5'- /5AmMC6/GGT GTG ATG TAT GTG GA/3ThioMC3-D/ -3'

Sequence 2 (16mer, 14 nm): 5'- /5ThioMC6-D/CCA CAT ACA TCA CAC C -3'

Sequence 3 (36mer, 25 nm): 5'- /5ThioMC6-D/TT TTT TTT TTT TTT TTT TTA CCA CAT ACA TCA CAC C -3'

Sequence 4 (76mer, 50 nm): 5'- /5ThioMC6-D/TT TTT TTT TTT TTT TTT TTT TTT TTT TTT TTT TTT TTT TTT TTT TTT TTT TTT TTT TTT ACC ACA TAC ATC ACA CC -3'

***Preparation of 3' fluorophore-labeled, 5' maleimide-functionalized DNA***

The 3' fluorophore-labeled, 5' maleimide-functionalized DNA oligonucleotide, to be conjugated to Fab' fragments, was synthesized as previously reported (1). Briefly, DNA oligonucleotide functionalized with the thiol group at 3'-end and the amino group at 5'-end (Sequence 1 above) was sequentially conjugated with the dye and the linker. First, ATTO647N-maleimide was conjugated to the thiol group at 3'-end, following manufacturer's instructions. Free dye was removed using a NAP-5 desalting column and ethanol precipitation. Labeling efficiency was verified using MALDI-TOF MS.  5'-end was then conjugated to the SM-$PEG_6$-maleimide linker by incubating with SM-$PEG_6$-maleimide in PBS (final concentration: 0.5 mM DNA-dye, 12.5-25 mM SM- $PEG_6$-maleimide, and 1x PBS). Half of the SM-$PEG_6$-maleimide was added first, and the remainder half was added after 30 min. The product was desalted over a NAP-5 column, precipitated in ethanol, and dried.

***Fab'-DNA synthesis***

In the following procedures, buffer exchange was performed using NAP-5 or PD-10 column and ultrafiltration was performed using Amicon Ultra spin filters, following the manufacturer's instructions.

The Armenian hamster anti-mouse CD3ε IgG 145-2C11 clone (2C11) and Armenian hamster anti-mouse TCRβ IgG H57-597 clone (H57) antibodies were fragmented, conjugated to DNA, and purified as previously reported (1). Because these two antibodies are the same isotype, the same protocol can be used for both. Briefly, H57 or 2C11, starting concentration > 3 mg/mL, was digested with pepsin at a pepsin to antibody ratio of 1:30 (by mass) for 8 h at 37 °C, with agitation, in 0.1 M acetate buffer pH 4.5, creating $F(ab')_2$ fragments. Digestion was quenched with 1/10 volume of 1 M Tris pH 8, then dialyzed in 1x PBS. Undigested IgG and Fc fragments were removed from the crude digest by incubating with protein A beads for 1 h, agitating, removing the supernatant, and washing the beads. Combined supernatant and washes were concentrated to > 0.5 mg/mL using spin filter. This $F(ab')_2$ preparation was then partially reduced to form Fab' fragments with 2 mM freshly prepared 2-mercaptoethylamine (2-MEA) in 2 mM EDTA, 1x PBS, incubating for 90 min. 2-MEA was removed over a desalting column. Maleimide DNA was immediately added in 2-20 fold molar excess and incubated at room temperature for 1-3 h, with agitation.

The rat anti-mouse CD3 IgG2b 17A2 clone (17A2) was fragmented using endoproteinase Glu-C from *Staphylococcus aureus* strain V8 (Glu-C) according to the previously reported procedure for Fab' purification from rat IgG2b antibodies with minor modifications (2). 17A2 was buffer-exchanged to 0.1 M phosphate buffer with 1 mM EDTA, pH 7.7 (1.99 mg/mL, 1260 µL). 3.3%w/w of Glu-C in the same buffer (1.53 mg/mL, 54.1 µL) was added and incubated at 37 °C, gently shaken for 17 h. The resulting mixture containing $F(ab')_2$ was buffer-exchanged to PBS with 2 mM EDTA and reacted with 2 mM 2-MEA for 90 min at 37 °C (protein concentration 0.66 mg/mL assuming IgG absorbance). 2-mercaptoethylamine was removed by buffer-exchange to the same buffer, yielding the mixture containing F(ab)' with reduced cysteine residues. The F(ab)' solution was concentrated with 30kDa MWCO spin filter (1.08 mL, protein concentration 0.96 mg/mL assuming IgG absorbance, corresponding to 13.8 nmol F(ab)') and reacted with 4.3 eq Atto647N-cBFL-maleimide (60 nmol in 100 µL) for 2 h at room temperature. Excess DNA was removed using repeated concentration-dilution using 30kDa MWCO spin filter. The crude product containing F(ab)'-DNA-Atto647N was purified by anion-exchange chromatography (Mono Q 5/50 GL from GE Healthcare, 200–1000 mM NaCl gradient in 20 mM Tris, pH 8.1), followed by size-exclusion chromatography (Superdex 75 Increase 10/300 from GE Healthcare, PBS). The obtained pure fractions were aliquoted and flash-frozen in liquid nitrogen with 10% glycerol and stored at -80 °C until being used. The reactions and purifications were monitored by non-reducing SDS-PAGE visualized by SYPRO Ruby staining (Invitrogen, Thermo Fisher) and the fluorescence from Atto647N.

***Thiol-DNA preparation***

Thiol DNA at a concentration of 1 mg/mL (197 µM) was reduced by treatment with 2 mM tris(2-carboxyethyl)phosphine (TCEP) in 10 mM HEPES pH 8 at 37 °C for 90 min. After incubation, the sample was desalted sequentially in two Bio-Spin 6 columns that had been equilibrated in PBS according to the manufacturer's direction. The concentration after desalting was measured using absorbance at 260 nm on a Nanodrop 2000 spectrophotometer.

***ICAM-1 and MHC class II I-E$^k$ preparation and peptide loading***
ICAM-1 with a decahistidin tag at its C-terminus (3) and MHC class II I-E$^k$ with hexahistidine tags on the C-terminus of both α and β-chains (4) were expressed and purified as previously described. Briefly, ICAM-1 (a gift from M. Davis) was packaged in baculovirus in SF9 cells, then transduced and expressed in High Five cells. MHC (a gift from L. Teyton and M. Davis) was expressed in S2 cells. For both constructs, expression was induced by $CuSO_4$ and the protein was purified by $Ni^{2+}$-NTA affinity column.

The MCC peptide (ANERADLIAYLKQATK) with a C-term GGSC linker was labeled with a Atto647N fluorophore with maleimide chemistry and purified by HPLC. On day 4, MCC-Atto647N was loaded into histidine-tagged MHC molecules by incubating in the loading buffer (PBS, pH adjusted to 4.5 with citric acid, 1% BSA). At the following day, loaded pMHC was isolated by filtering unloaded MCC peptide with 10 kDa MWCO Amicon Ultra spin filter.

***Supported lipid bilayer (SLB) preparation***
SLBs were formed as reported previously in Attofluor cell chambers (Invitrogen, Thermo Fisher). Number 1.5 25 mm round coverslips (Warner Instruments, Holliston, MA, USA) were ultrasonicated in 50:50 isopropanol:water for 30 min, then rinsed thoroughly in Milli-Q water (EMD Millipore, Billerica, MA, USA). Cleaned coverslips were then etched for 3-5 min in piranha solution (3:1 sulfuric acid:hydrogen peroxide), and again rinsed thoroughly in Milli-Q water.

Vesicles for bilayer formation were prepared by first mixing 95% DOPC, 2% Ni-NTA DGS, and 3% MCC-DOPE phospholipids, by mol, in chloroform, drying on a rotovap, and then resuspending to 0.5 mg/mL in water. Small unilamellar vesicles (SUVs) were prepared by probe sonication for 1 min total with pulses of 15 s on, 10 s off. The lipid solution was kept in an ice bath to prevent the temperature from rising during the sonication. Sonicated solutions were then centrifuged at 21,000 × g for 20 min at 4 °C to remove lipid aggregates and Ti particles. Vesicle solutions in water were mixed 1:1 with 1x PBS, added to imaging chambers (300 µL per chamber), and incubated for 30 minutes. The SLBs were then rinsed and thiol-DNA (seq. 2-4, reduced and desalted as described in Thiol-DNA preparation) was injected at a final concentration of 1 µM and incubated for 80-120 minutes. Bilayers were stable overnight at 4 °C.

The membrane was then incubated with 100 mM $NiCl_2$ in Tris-buffered saline (TBS) for 5 min, rinsed with TBS, and then rinsed with imaging buffer (1 mM $CaCl_2$, 2 mM $MgCl_2$, 20 mM HEPES, 137 mM NaCl, 5 mM KCl, 0.7 mM $Na_2HPO_4$, 6 mM D-glucose, and 0.2% w/v bovine serum albumin (BSA)). Addition of BSA blocks any defects in the bilayer to ensure that proteins of interest cannot stick non-specifically to the surface and instead conjugate specifically to the mobile lipid bilayer. BSA was added to bilayers no more than 2 h before imaging to ensure quality bilayers. A solution containing all proteins to be coupled to the membrane (Fab'-DNA and/or MCC pMHC and ICAM-1) was then prepared in imaging buffer, added to the sample, incubated for 30-35 min, and then washed thoroughly with imaging buffer. All SLBs were equilibrated for 15 min to 37 °C before adding cells. All imaging was done at 37 °C.

***T-cell harvesting and culturing***
CD4+ T cells expressing the AND TCR were harvested, cultured, and transduced as previously described (5, 6). Briefly, T cells were harvested (day 1) from the cross of (B10.Cg-Tg(TcrAND)53Hed/J) x (B10.BR-H2k2 H2- T18a/SgSnJ) strains transgenic mice (The Jackson Laboratory, Bar Harbor, ME, USA) and activated by 2 µM moth cytochrome *c* peptide, amino acids 88−103, (ANERADLIAYLKQATK) (MCC) in RVC media immediately after harvest. IL-2 was added 24 hours after harvest (day 2). On day 3, activated T cells were retrovirally

transduced with LAT-eGFP or NFAT-mCherry-containing supernatants (MSCV vector) collected from Platinum-Eco cells (Cell Biolabs, San Diego, CA, USA) in RVC media. On day 4, T cell media was exchanged with fresh RVC and IL-2 to remove viral particles. T cells were imaged on days 5 to 8, changing media regularly to maintain proper nutrients and pH. All animal work was approved by Lawrence Berkeley National Laboratory Animal Welfare and Research Committee under the approved protocol #17702.

***Microscopy***

*Equipment*

Total internal reflection fluorescence (TIRF) microscopy experiments were performed on a motorized inverted microscope (Nikon Eclipse Ti-E; Technical Instruments, Burlingame, CA, USA) equipped with a motorized Lumen Dynamics X-Cite® 120LED Fluorescence Illumination System Epi/TIRF illuminator, (Excelitas Technologies, Waltham, MA, USA), Perfect Focus system, and a motorized stage (Applied Scientific Instrumentation MS-2000, Eugene, OR, USA). A laser launch with 488, 561, and 640 nm (Coherent OBIS, Santa Clara, CA) diode lasers was controlled by an OBIS Scientific Remote (Coherent Inc., Santa Clara, CA) and aligned into a fiber launch custom built by Solamere Technology Group, Inc. (Salt Lake City, UT, USA). A dichroic beamsplitter (z488/647rpc; Chroma Technology Corp., Bellows Falls, VT, USA) reflected the laser light through the objective lens and fluorescence images were recorded using an EM-CCD (iXon 897DU; Andor Inc., South Windsor, CT, USA) after passing through a laser-blocking filter (Z488/647M; Chroma Technology Corp., Bellows Falls, VT, USA). Exposure times, multidimensional acquisitions, and time-lapse periods for all experiments were set using Micro-Manager (7). A TTL signal from the appropriate laser triggered the camera exposure.

*Image acquisitions*

The laser intensities were measured at the sample for each experiment day, so that constant laser intensities are used for each type of imaging across different days. Imaging of Fab'-DNA diffusion on the supported membrane was performed with a streaming acquisition of 14 ms at 8.6 mW power at the sample for Fab'-DNA-Atto647N.

To gather step photobleaching statistics for each Fab'-DNA construct, gel-phase bilayers presenting Fab'-DNA were formed by substituting the DOPC used in fluid bilayers for DPPC. In fluid bilayers, the rapidly diffusing particles made it difficult to track a single particle throughout its trajectory until bleaching, as particles frequently crossed paths or diffused out of the field of view. On gel-phase bilayers, particles were essentially immobile over the ~2.4 s acquisition taken to assess the number of photobleaching steps from each particle. For both 2C11 and 17A2 Fab'-DNA constructs >99% of fluorescent signals bleached in a single step (221/222 ≈ 99.5% for 2C11 Fab'-DNA and 273/275 ≈ 99.3% for 17A2) (Figs. 2e,g). The low-probability two-step photobleaching events are most likely from two particles stochastically adhered to the gel-phase SLB at a distance below the diffraction limit, which would occur with a probability of 0.43% on bilayers with a particle density of 0.07 $\mu m^{-2}$, agreeing extremely well with the experimental data.

For Fab'-DNA dwell time measurements, TIRF images of long exposure time (500 ms) using a laser power of 0.8 mW (640 nm) at the sample were collected every 10 s. Photobleaching data were taken for each species using the same laser power and exposure time as was used to collect dwell time data, but without a time lapse in order to minimize recovery after photobleaching. The fraction bound measurements were taken with power at the sample of 4.3 mW (640 nm), exposure times of 20 ms and 500 ms, and EM gains of 1000 and 50 to resolve all and bound Fab'-DNA molecules, respectively. Images for all and bound ligands were taken sequentially. Cell footprint for both of these measurements was determined by RICM.

For NFAT titrations, cells were either transduced with NFAT-mCherry or with LAT-eGFP P2A NFAT-mCherry. Transduced cells were identified in TIRF in either the mCherry or eGFP channel, depending on constructs used, such that activation state could not be assessed. Then RICM, binding event (TIR 640), LAT (TIR 488) and NFAT (epi 561) snapshots were taken of the cell. NFAT snapshots were taken at 0, 3, and 6 μm above the bilayer in order to clearly identify the nucleus. T cell sensitivity to ligand was identical for the P2A and NFAT-only constructs. The ligand density for 17A2 Fab'-DNA was modified to be 70% of the measured fluorescence density on the bilayer due to the contamination of DNA-dye species conjugated to smaller antibody fragments (Fig. S2).

To investigate LAT condensation in response to single binding events, long exposure (500 ms), low power (0.8 mW) TIR 640 images of binding events were immediately followed by TIR 488 images of LAT at 50-200 ms, 0.4-1 mW power, with precise imaging conditions depending on the LAT expression level of the cell in order to image with the best dynamic range. Image stacks were acquired with a 2 s time lapse.

### ***Image Analysis***

*Diffusion Analysis*
Fluorescent particles were tracked using the FIJI plugin TrackMate (8). Particles were identified using the difference of Gaussians detector, and diameters, intensity thresholds, and maximum linking distances were set by eye and then all data were uniformly analyzed. The particle density in movies was very low (0.01 $\mu m^{-2}$) to accurately track single particles for 10's to 100's of frames. In the rare occasion that two particles overlapped, only one particle was localized and this localization was used to populate the step size distribution along one track. The other particle was split into a second track. Because overlapping events are very rare at these densities, this is expected to have negligible impact on the final step size distribution. The particle localization and linking data were exported to MATLAB using a custom python script. A custom MATLAB script was then used to build step size distributions as a function of time delay between frames. The step size distribution between particles in adjacent frames (14 ms) contained artifacts because the step sizes were too small compared to the camera pixel size (0.107 μm) and the algorithm's localization error. This time delay was therefore removed from analysis. For subsequent time delays, distances were calculated between every other frame, every third frame, etc. drawing from the same tracking data. Skips were used rather than a scanning window to prevent over-counting. The entire data set was then fit to a single component diffusion model (Eq. 1 in the main text) to extract the diffusion coefficient, $D$. Error was calculated using a 95% confidence interval.

*Step photobleaching*
Particles were tracked as described above. Because step photobleaching data were taken on DPPC bilayers, the maximum linking distance allowed was very small, 0.2 μm. After particle photobleaching, the mean background at the particle's last visible location was measured for 25 subsequent frames. The intensity trace from the beginning of the acquisition through the 25 frames after bleaching were fit were analyzed using a Bayesian change point detection algorithm with a MATLAB script. As described above, one trace for 2C11 Fab'-DNA and two traces for 17A2 Fab'-DNA were identified to undergo two photobleaching steps. This corresponds to the probability that two particles were randomly attached to the gel-phase bilayer at a distance below the diffraction limit.

*Dwell time*
Binding events were tracked using TrackMate. Particle intensity and linking distances were first applied automatically. After initial cell landing and spreading, bound ligand undergoes directed motion to the geometric center of the cell. TrackMate is not well-designed for this type of motion, and so tracks were manually edited to reflect the most likely true tracks based on this additional information about the system. The long dwell time of Fab'-DNA complicates accurate tracking of binding events for their full duration. Binding events may overlap in the periphery or become difficult to track as they all convene in the kinapse, which usually occurs within two minutes of peripheral ligand binding. This leads to a systematic undercounting of long Fab'-DNA:TCR dwell times and an excess of short, artificially cut-off trajectories. Fluorescent spots resolved for only one frame were discarded from analysis due to occasional spurious localization errors. The dwell time distribution was built and fit to a single exponential decay using a custom MATLAB script.

The photobleaching data for each ligand were analyzed by cropping the images to a region of even illumination, background subtracting, and calculating the mean intensity of each frame. The mean intensity was then plotted as a function of time and fit to a double exponential decay due to a dim, quickly bleaching background contaminant. The longer time constant was then multiplied by the time lapse used in acquiring the dwell time data to accurately capture the photobleaching rate. The bleaching rate is subtracted from the observed off-rate to determine the mean single molecule dwell times.

*Fraction bound*
The RICM channel was used to mask images of all and bound Fab'-DNA molecules below single cells. The mask was created by thresholding the RICM image using isodata thresholding, removing small objects and objects touching the edge, filling holes, and then slightly dilating the identified area to more accurately capture membrane ruffles and filopodia at the SLB – T cell interface. Particles were localized using TrackMate and number of bound Fab'-DNA was divided by the total number of Fab'-DNA for each cell to measure the fraction bound. This fraction may exceed 1 if a Fab'-DNA molecule bound TCR during the long-exposure image, which was taken after the corresponding short-exposure image.

*NFAT localization*
The cytosol and nucleus of T cells were segmented using the ilastik pixel classification workflow (9). Images of NFAT-mCherry were used to train the classifier to identify pixels belonging to the cytsol and nucleus of a cell. Reliable segmentation was accomplished with the following selected image features: Color/Intensity: $\sigma = 0.7$ pixels; Edge: $\sigma = 1$ and $\sigma = 3.5$; Texture: $\sigma = 3.5$. Use of the edge and texture features allowed segmentation regardless of whether the NFAT-mCherry signal was greatest in the cytosol or the nucleus. All images were manually inspected for correct segmentation. The background-subtracted intensities in each region were then calculated, and.a nucleus:cytosol intensity ratio greater than 1 indicated activation. Only cells with substantial interfacial contact areas (based on RICM signal) and with unambiguous nuclei were included in the quantification of endpoint NFAT activation state.

*LAT condensate analysis*
Binding events were tracked as described for the dwell time analysis. These data were taken with a 2 s, rather than 10 s time lapse, and so while fluorophores bleached more quickly, they were also more easily tracked.

LAT condensates were first identified using the ilastik pixel classification workflow (9). First, time series of LAT images from cells with four different background LAT intensities were loaded as input data so that the algorithm could accurately identify LAT condensates in cells with low, medium, and high LAT expression levels. Reliable identification of LAT condensates was accomplished with the following selected image features – Color/Intensity: $\sigma$ = 0.7; Edge: $\sigma$ = 1.6; Texture: $\sigma$ = 0.7 and $\sigma$ = 5. The pixel classifier was then trained to identify image background, cell background, and LAT condensates. The success of the algorithm was assessed on the four training stacks and four non-training stacks. Once LAT pixels appeared to be identified reliably, all data were processed using the same pixel classifier.

Pixel probabilities were than exported to FIJI, where LAT pixel probabilities were tracked in TrackMate. LAT condensates were identified with the Laplacian of Gaussian (LoG) detector, with the particle diameter set to 7 pixels and the threshold set to 1000. These parameters worked well across LAT condensate sizes and cells, thanks to accurate pixel probabilities produced by ilastik. LAT condensates were then tracked using a maximum linking distance of 7 pixels and allowing gaps of 1 frame. Merging and splitting were not allowed. Tracks were counted as LAT condensates if they persisted for at least 4 frames. This tracking method was robust and required little manual editing.

Binding event and LAT data were then exported to MATLAB using a custom python script, where the ratio of LAT condensates to binding events was calculated and plotted for each cell.

**Statistical Analysis and Reproducibility**
Measurements of Fab'-DNA diffusing freely on the SLB were replicated on at least three independently prepared bilayers and data shown are representative. All conditions for all live cell measurements (Figs 3-5) were replicated with cells from at least two mice, and in many cases three mice. Dwell time distributions, fraction bound, and $N_{LAT}/N_{binding\ events}$ show compiled data for all cells analyzed. NFAT translocation dose-response curves are representative of at least two experiments. The Mann-Whitney U-test was used to determine if distributions were significantly different for Fig. 5c. p-values for pairwise comparisons and number of cells per condition are listed in Supplementary Tables 1 and 2.

Supporting References


1. Lin, J.J., G.P. O'Donoghue, K.B. Wilhelm, M.P. Coyle, S.T. Low-Nam, N.C. Fay, K.N. Alfieri, and J.T. Groves. 2020. Membrane Association Transforms an Inert Anti-TCRβ Fab' Ligand into a Potent T Cell Receptor Agonist. *Biophys. J.* 118:2879–2893.
2. Rousseaux, J., R. Rousseaux-Prévost, H. Bazin, and G. Biserte. 1983. Proteolysis of rat IgG subclasses by Staphylococcus aureus V8 proteinase. *Biochim. Biophys. Acta - Protein Struct. Mol. Enzymol.* 748:205–212.
3. Hartman, N.C., J.A. Nye, and J.T. Groves. 2009. Cluster size regulates protein sorting in the immunological synapse. *Proc. Natl. Acad. Sci. U. S. A.* 106:12729–12734.
4. Nye, J.A., and J.T. Groves. 2008. Kinetic control of histidine-tagged protein surface density on supported lipid bilayers. *Langmuir*. 24:4145–4149.
5. Smith, A.W., A.A. Smoligovets, and J.T. Groves. 2011. Patterned Two-Photon Photoactivation Illuminates Spatial Reorganization in Live Cells. *J. Phys. Chem. A*. 115:3867–3875.
6. O 'Donoghue, G.P., R.M. Pielak, A.A. Smoligovets, J.J. Lin, and J.T. Groves. 2013. Direct single molecule measurement of TCR triggering by agonist pMHC in living primary T cells. *Elife*. 2.

7. Edelstein, A., N. Amodaj, K. Hoover, R. Vale, and N. Stuurman. 2010. Computer Control of Microscopes Using µManager. *Curr. Protoc. Mol. Biol.* 92.
8. Tinevez, J.Y., N. Perry, J. Schindelin, G.M. Hoopes, G.D. Reynolds, E. Laplantine, S.Y. Bednarek, S.L. Shorte, and K.W. Eliceiri. 2017. TrackMate: An open and extensible platform for single-particle tracking. *Methods*. 115:80–90.
9. Berg, S., D. Kutra, T. Kroeger, C.N. Straehle, B.X. Kausler, C. Haubold, M. Schiegg, J. Ales, T. Beier, M. Rudy, K. Eren, J.I. Cervantes, B. Xu, F. Beuttenmueller, A. Wolny, C. Zhang, U. Koethe, F.A. Hamprecht, and A. Kreshuk. 2019. Ilastik: Interactive Machine Learning for (Bio)Image Analysis. *Nat. Methods*. 16:1226–1232.

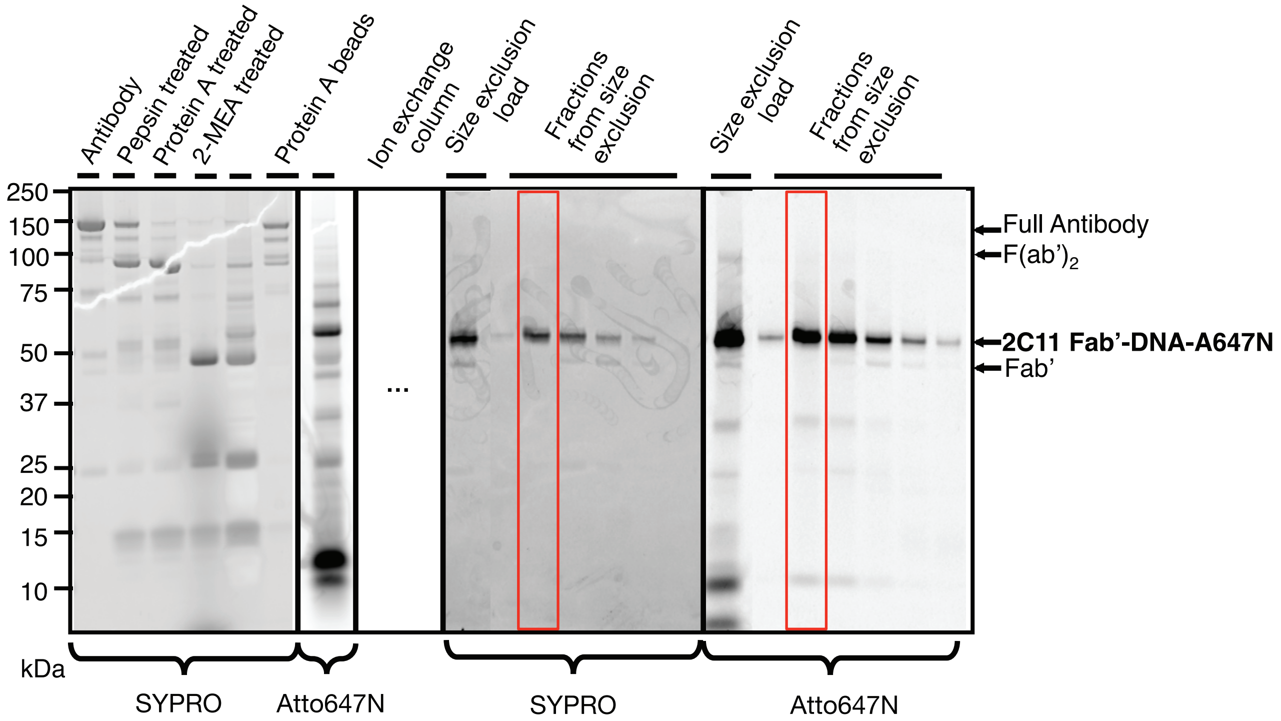


Supporting Figure 1. The fragmentation of 2C11 is monitored by SDS-PAGE and the purity of 2C11 Fab'-DNA-Atto647N after purification over ion exchange and size excusion columns is confirmed.

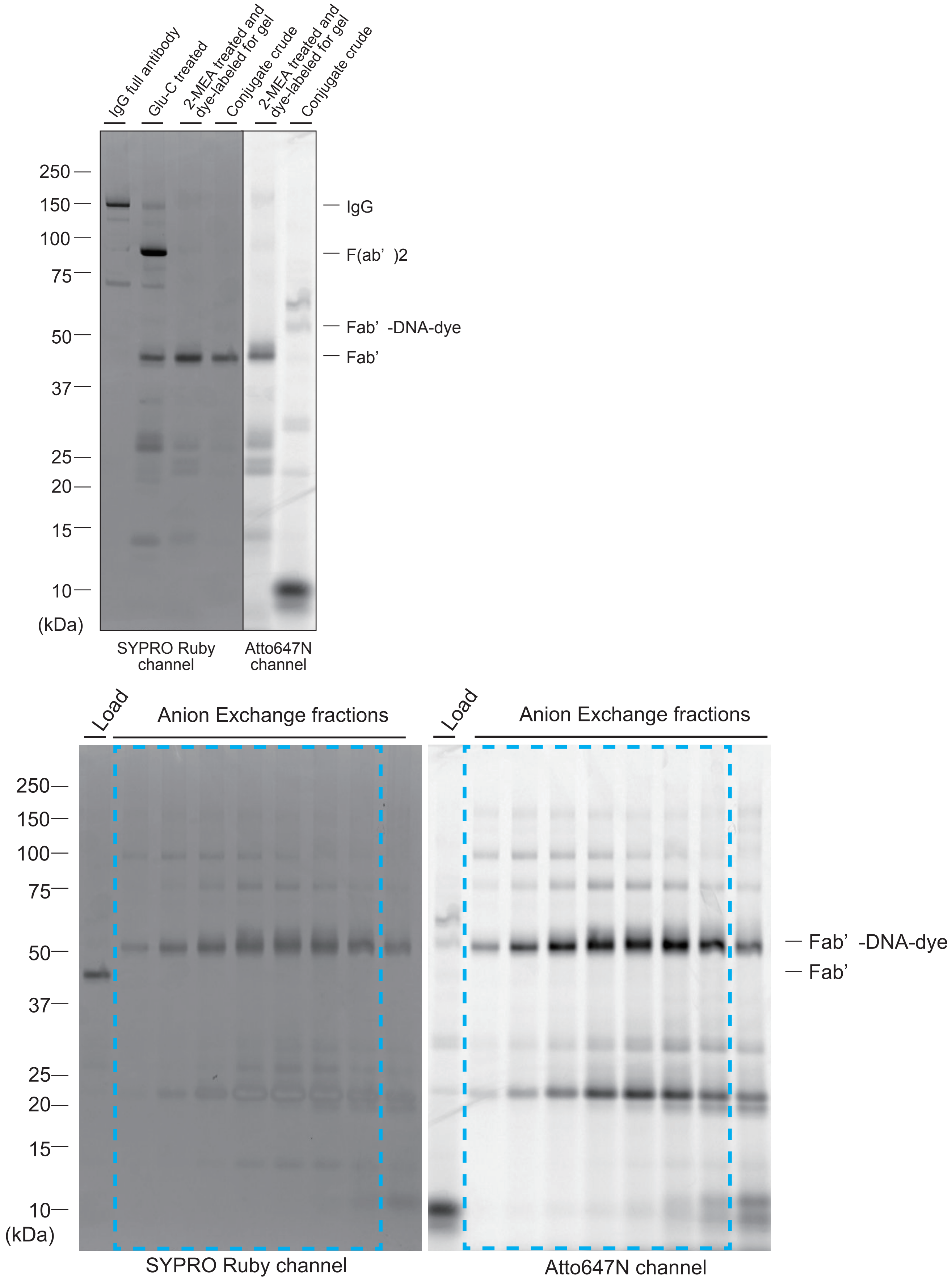
IgG full antibody
Glu-C treated
2-MEA treated and dye-labeled for gel
Conjugate crude
2-MEA treated and dye-labeled for gel
Conjugate crude
250
150
100
75
50
37
25
20
15
10
(kDa)
IgG
F(ab’ )2
Fab’ -DNA-dye
Fab’
SYPRO Ruby channel
Atto647N channel
Load
Anion Exchange fractions
Load
Anion Exchange fractions
250
150
100
75
50
37
25
20
15
10
(kDa)
Fab’ -DNA-dye
Fab’
SYPRO Ruby channel
Atto647N channel

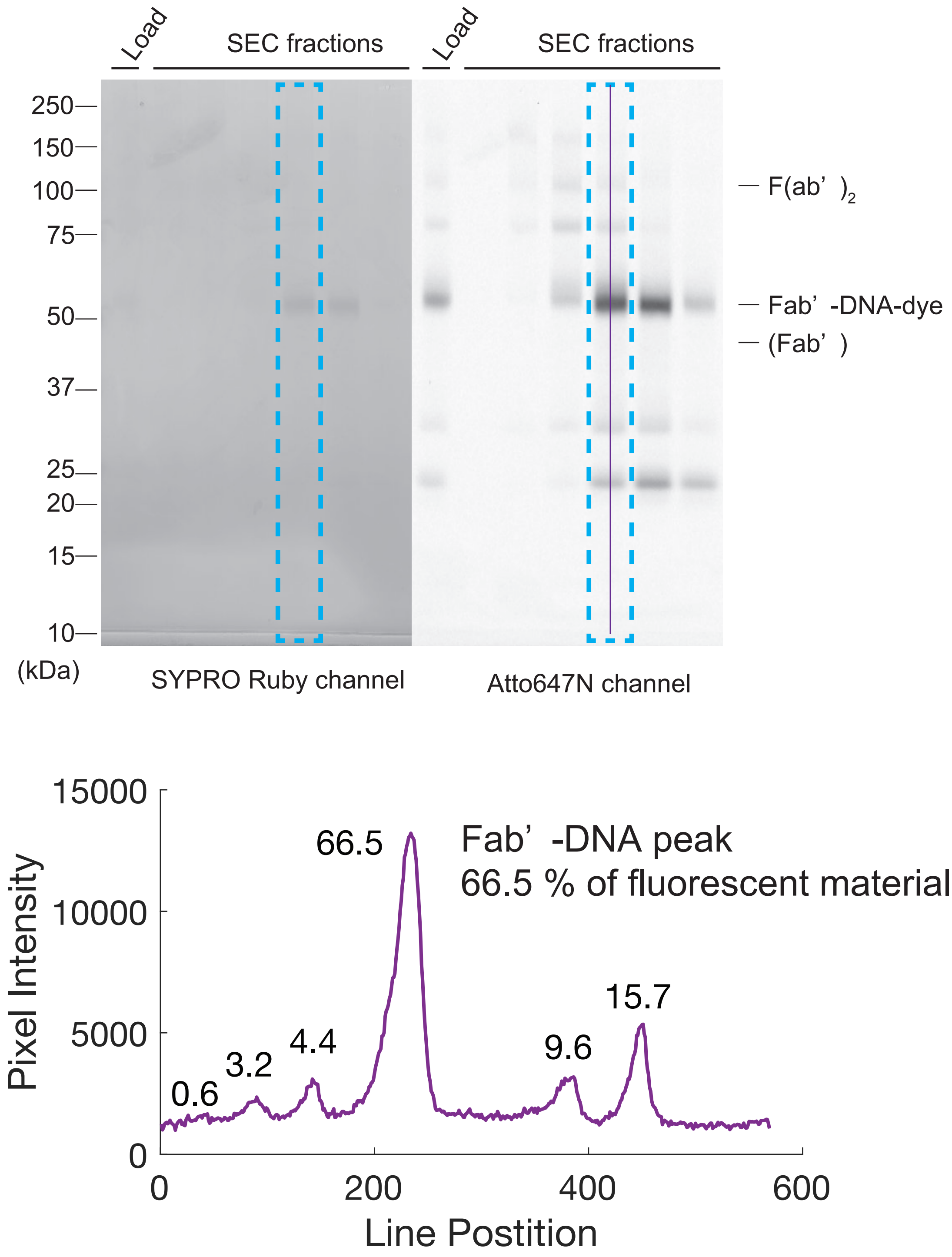


Supporting Figure 2. The fragmentation of 17A2 is monitored by SDS-PAGE and the purity of 17A2 Fab'-DNA-Atto647N after purification over ion exchange and size excusion columns is assessed. A line scane of the purified Fab'-DNA shows that about two thirds of the fluorescent species are Fab'-DNA and the majority of the contaminants are smaller DNA-conjugated fragments which cannot bind TCR.

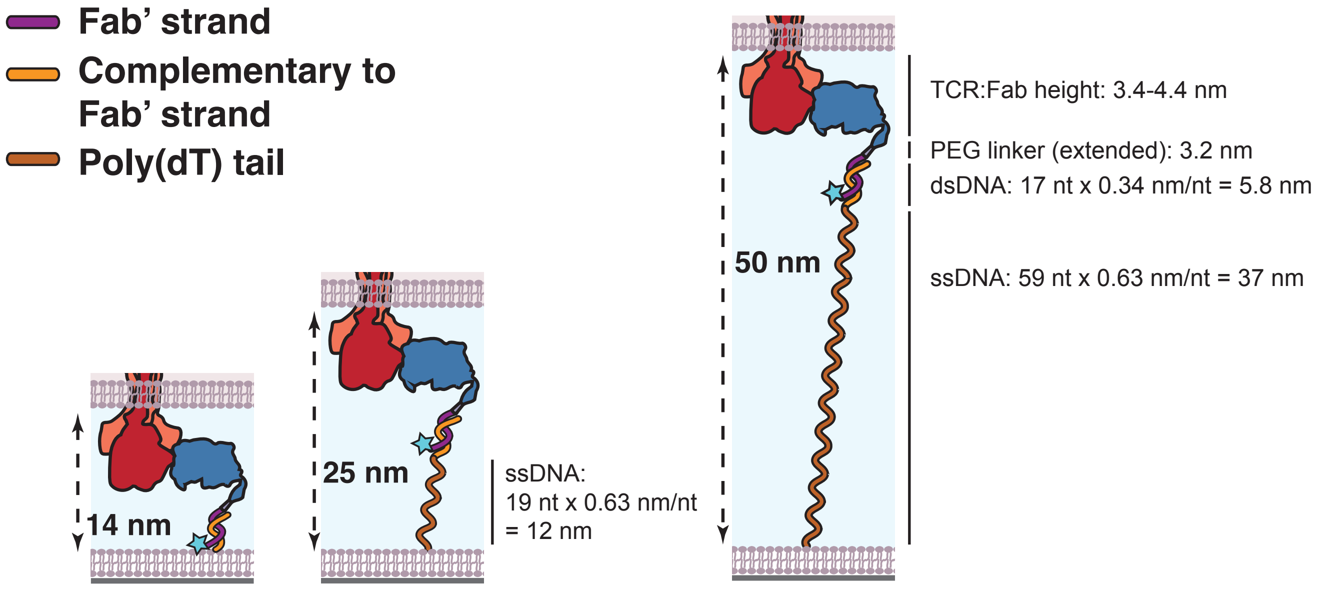


Supporting Figure 3. Approximation of intermembrane space allowed at binding events between Fab’-DNA and TCR, assuming oligonucleotides can fully stretch if needed. The inter-membrane space for each thiol-DNA tether was estimated using the structure of H57 Fab bound to TCR (PDB: 1NFD), the estimated length of the PEG linker, the height of a double stranded DNA base pair, and the height of a single stranded DNA nucleotide.

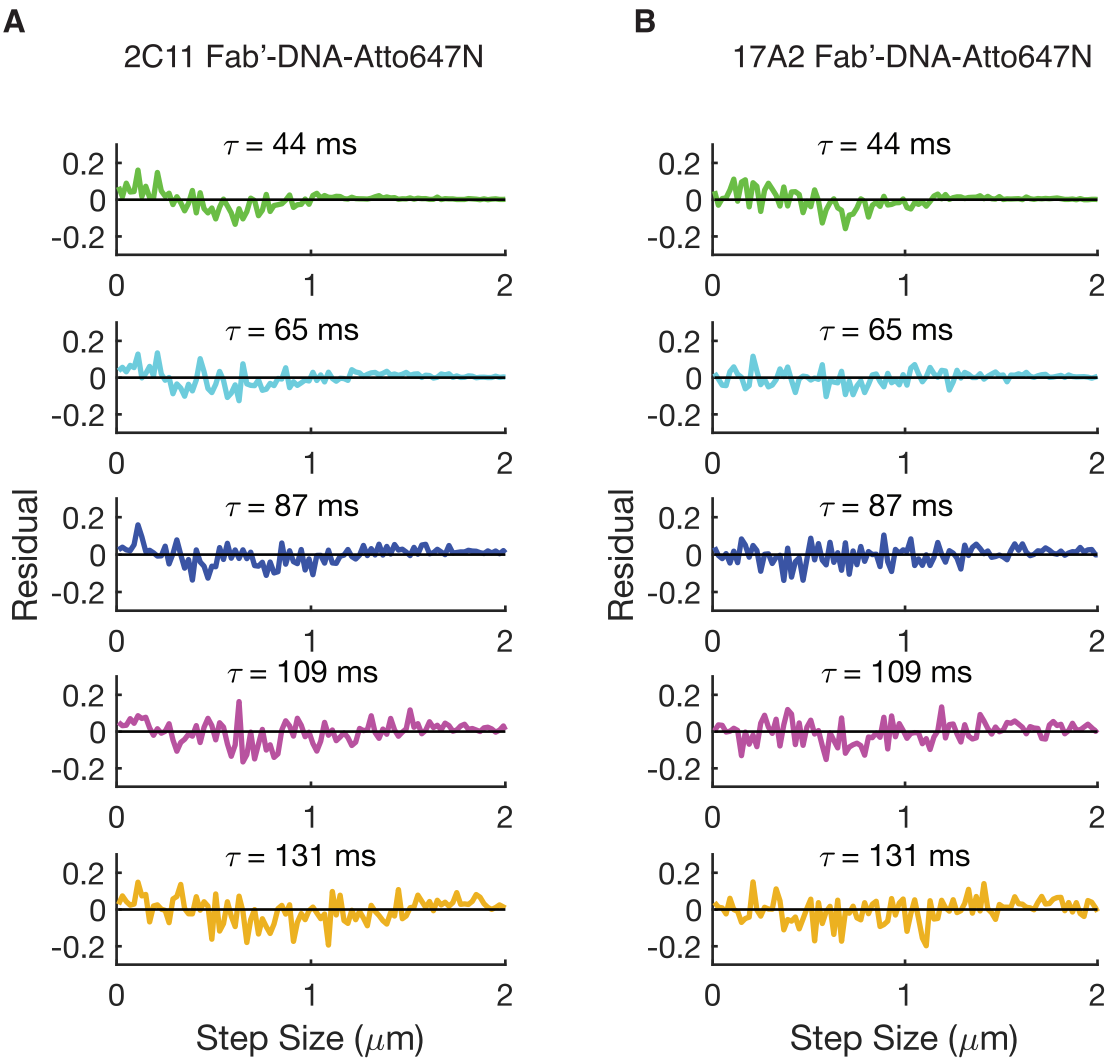


Supplementary Figure 4. Residuals from **(A)** 2C11 Fab'-DNA and **(B)** 17A2 Fab'-DNA step size distribtuions at multiple delay times, fit by a two-dimensional single component Brownian diffusion model. Data from all time delays were fit simultaneously with eqn. 1 to obtain a single diffusion coefficient for each Fab'-DNA construct.

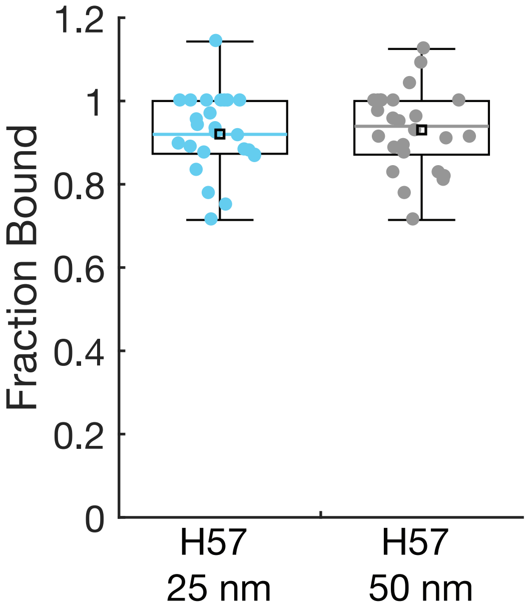


Supporting Figure 5. The fractions of H57 Fab'-DNA with medium and long tether lengths that are bound under T cells are nearly identical to each other and to H57 Fab'-DNA with the short tether. Colored bar: median; black square: mean; box: interquartile range; whiskers: data within 1.5x IQR. H57 25 nm: n = 25; H57 50 nm: n = 26.

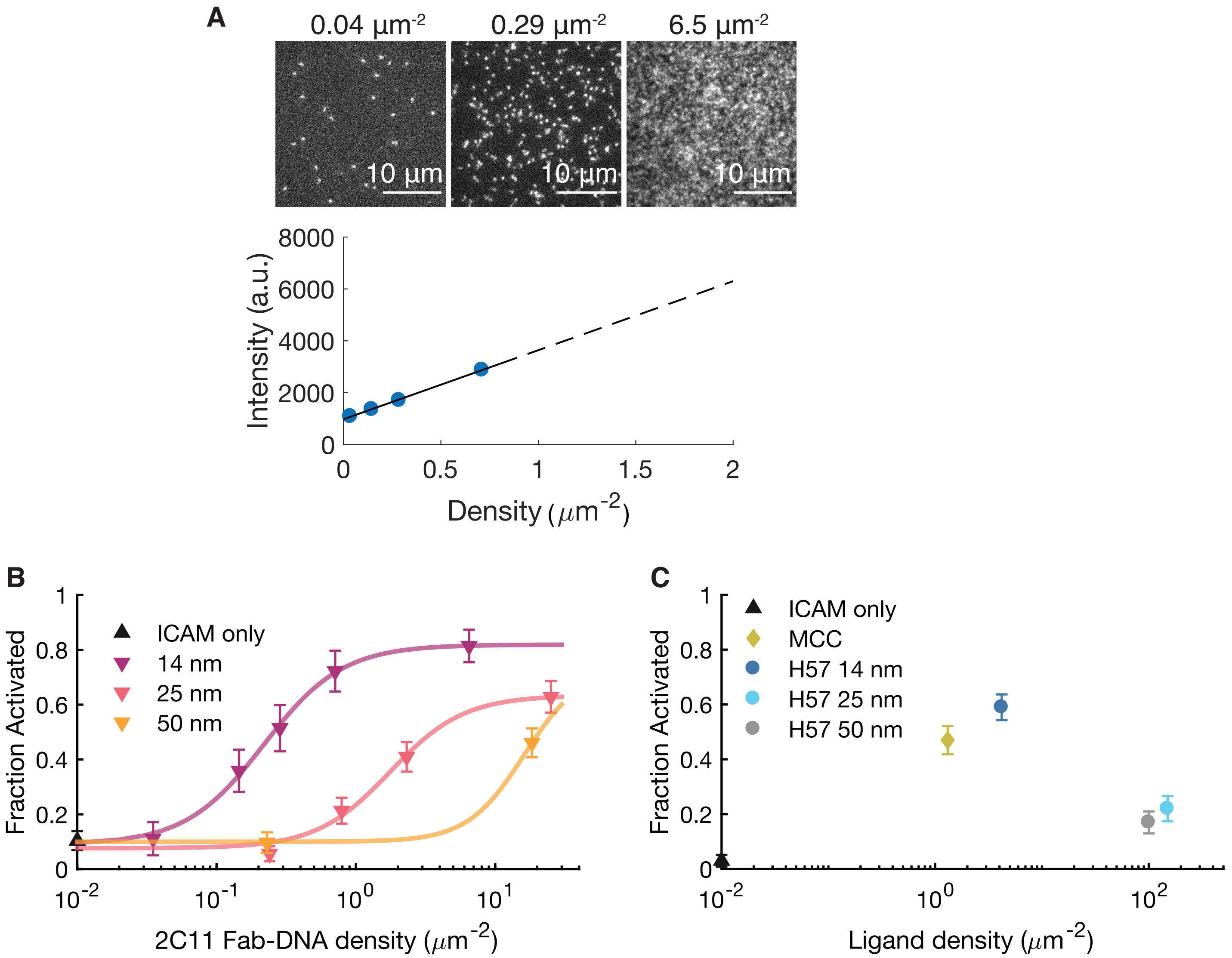


Supporting Figure 6. (**A**) The density of ligand on the bilayer is determine by TIRF intensity. The total intensity and particle density are measured at densities for which particles are countable. The density vs. fluorescence intensity calibration curve is then extrapolated to determine the density of ligand on high-density bilayers. (**B**) NFAT titration curves for 2C11 Fab'-DNA with varied tether lengths. The inflection points for all three constructs roughly match those for the corresponding H57 constructs. **(C)** Even at very high (~100 $\mu m^{-2)}$ density, H57 Fab'-DNA constructs that allow up to 25 nm and 50 nm intermembrane space do not fully activate T cells compared to the 14 nm Fab'-DNA and pMHC controls, though they do activate significantly above the ICAM only negative control. Of note, cells in the experiment shown had a low maximal fraction of cells that activated in response to short ligands (~0.6 compared to ~0.8), which may relate to why the fraction activated, especially from the medium tether ligand, is low compared to the data set shown in panel (B) and Fig. 4D.

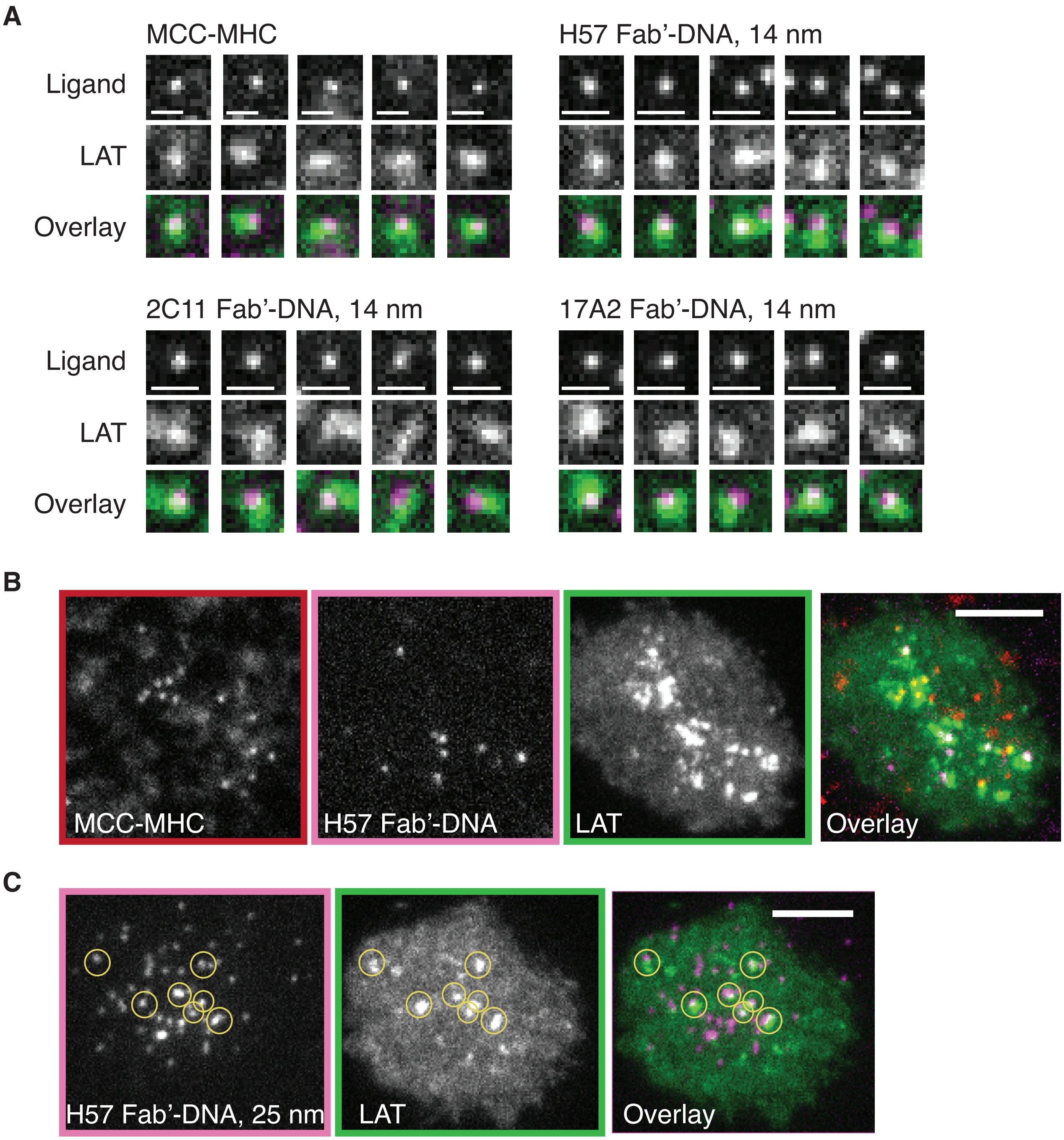


Supporting Figure 7. LAT clustering in response to binding events. (**A**) Examples are shown of single binding events colocalized with LAT clusters for potent ligands. Scale bars 1 μm. (**B**) LAT condensates form in response to binding events between TCR and both H57 Fab'-DNA-AlexaFluor555 (14 nm) and pMHC-Atto647N when both ligands are presented at low density on the SLB. Binding events from both ligands appear to have an additive effect on signal tranduction. Scale bar 5 μm. (**C**) Instances where LAT clusters form in response to weak ligands often colocalize with a cluster of binding events. Scale bar 5 μm.

Supplementary Table 1. Mann-Whitney U Test statistics for pairwise comparisons of $N_{LAT}/N_{bind}$ distributions in Fig. 5b.

| | | p | zval | ranksum |
|---|---|---|---|---|
| **Comparing epitope (all 14 nm)** | | | | |
| **ligand 1** | **ligand 2** | | | |
| MCC | 2C11 | 0.45 | -0.75 | 143 |
| MCC | 17A2 | 0.027 | -2.22 | 95.5 |
| MCC | H57 | 0.37 | -0.89 | 140 |
| 2C11 | 17A2 | 0.049 | -1.97 | 210 |
| 2C11 | H57 | 0.73 | -0.34 | 287 |
| 17A2 | H57 | 0.06 | 1.79 | 221 |
| **Comparing tether length** | | | | |
| **ligand 1** | **ligand 2** | | | |
| H57 14 nm | H57 25 nm | 0.00029 | 3.62 | 390 |
| H57 14 nm | H57 50 nm | 0.000042 | 4.09 | 320 |
| H57 25 nm | H57 50 nm | 0.178 | 1.35 | 242 |

Supplementary Table 2. Number of cells analyzed for each ligand condition in Fig. 5b. Cells were only analyzed if they experienced at least 20 binding events to ensure reasonable statistics for each cell.

| ligand | n cells analyzed |
|---|---|
| MCC 14nm | 11 |
| 2C11 14 nm | 17 |
| 17A2 14 nm | 12 |
| H57 14 nm | 17 |
| H57 25 nm | 16 |
| H57 50 nm | 10 |

Movie S1. 2C11 Fab'-DNA freely diffuses in two dimensions on a supported lipid bilayer. 2C11 Fab'-DNA density is 0.09 $\mu m^{-2}$. Scale bar 5 μm.

Movie S2. 17A2 Fab'-DNA freely diffuses in two dimensions on a supported lipid bilayer. 17A2 Fab'-DNA density is 0.06 $\mu m^{-2}$. Scale bar 5 μm.

Movie S3. T cells land and spread on the bilayer, visualized with RICM (left), and bind Fab'-DNA, visualized by TIRF (right). Fab'-DNA stays bound for 10's of seconds as bound TCR are tracked to the geometric center of the cell. Scale bar 5 μm.
